# *Dnmt3a*-mutant Leukemia Stem Cells evade chemotherapy through enforced quiescence in *Npm1^c^-Flt3^ITD^* Acute Myeloid Leukemia

**DOI:** 10.64898/2026.06.03.729757

**Authors:** Paniz Tavakoli Shirazi, Jasmin Straube, Victoria Ling, Stacey Andersen, Emily Cooper, Siu Hang Natalie Chan, Rohit Haldar, Yashaswini Janardhanan, Leanne Cooper, Claudia Bruedigam, Carolyn Grove, Megan Bywater, Steven W Lane

**Affiliations:** Cancer Research Program, QIMR Berghofer Medical Research Institute, Brisbane, Australia; The University of Queensland, St Lucia, Brisbane, QLD, Australia; The Australian Centre for Blood Diseases, School of Translational Medicine, Faculty of Medicine, Nursing and Health Sciences, Monash University, Melbourne, VIC; University of Western Australia, School of Pathology and Laboratory Medicine, Perth, WA, Australia; Cancer Care Services, Royal Brisbane and Women’s Hospital, Brisbane, QLD, Australia

## Abstract

Concurrent mutations in *DNMT3A, NPM1*, and *FLT3* define a high-risk subtype of acute myeloid leukemia (AML) associated with increased relapse risk and inferior survival following standard chemotherapy. However, the mechanisms by which *DNMT3A* mutations promote treatment resistance in *NPM1*^*c*^*-FLT3*^*ITD*^ AML remain unclear.

Using genetically engineered murine models of *Npm1*^*c*^*-Flt3*^*ITD*^ AML with or without *Dnmt3a*^*R878H*^ (homologous to human *DNMT3A*^*R882H*^), we demonstrate that *Dnmt3a*^*R878H*^ promotes chemotherapy resistance through epigenetic regulation of leukemia stem cell (LSC) quiescence. Integrated transcriptomic and epigenetic profiling revealed coordinated remodeling of DNA methylation and chromatin accessibility in LSC-enriched populations, characterized by preferential hypomethylation and increased accessibility at loci associated with stemness and quiescence programs. These data were confirmed in human *DNMT3A*^*R882H*^*-NPM1*^*c*^*-FLT3*^*ITD*^ AML datasets with enrichment of quiescence-associated and stem cell enriched transcriptional programs. Conversely, *Dnmt3a*-mutant LSCs retained sensitivity to the cell-cycle independent regimen venetoclax plus azacitidine, but residual LSCs exhibited transcriptional plasticity and reversion to a de-differentiated state.

We have identified LSC heterogeneity spanning primitive hematopoietic stem cell (HSC)-and progenitor-like states and our data demonstrate preferential maintenance of a quiescent HSC-like LSC subpopulation in *Dnmt3a*^*R878H*^-mutant AML following chemotherapy treatment.

Pharmacologic induction of cell-cycle entry using pegylated interferon α (pegIFNα) disrupted the quiescent LSC state and restored chemotherapy sensitivity, identifying quiescence as a reversible and therapeutically actionable mechanism of resistance.

These findings identify *DNMT3A*-mediated epigenetic regulation of LSC quiescence as a conserved mechanism of standard chemotherapy resistance and position therapeutic reactivation of quiescent LSCs as a promising strategy to overcome chemotherapy resistance and improve outcomes in high-risk *DNMT3A*-mutant AML.

## Introduction

Patients with acute myeloid leukemia (AML) who experience relapse following intensive chemotherapy or low-intensity venetoclax combinations have an extremely adverse prognosis ^1-3^. The molecular and cytogenetic profile of AML at diagnosis is a powerful predictor of relapse and overall survival after treatment ^4-6^.

Mutations in nucleophosmin 1 (*NPM1*) are the most common recurrent molecular abnormality in *de novo* AML, frequently co-occurring with activating mutations in Fms-like tyrosine kinase 3, *FLT3* (∼60%) and DNA methyltransferase 3A, *DNMT3A* (∼50%) ^5^. Patients harboring the triple combination of *DNMT3A*^*mut*^*-NPM1*^*c*^*-FLT3*^*ITD*^ account for ∼10% of all AML cases and ∼25% of *NPM1*^*c*^*-*mutated AMLs ^4,5^. We and others have shown that the combination of these three mutations has a major negative impact on prognosis ^7-13^. Functionally, these mutations cooperate to accelerate leukemia initiation and progression *in vivo* ^14-16^. *DNMT3A* mutations in *NPM1*^*c*^ AML are associated with higher rates of measurable residual disease (MRD) post-chemotherapy, an indicator of high relapse-risk ^12,17^. These observations define *DNMT3A*^*mut*^*-NPM1*^*c*^*-FLT3*^*ITD*^ as an aggressive, chemotherapy-resistant AML subtype, however the mechanisms underlying *DNMT3A*^mut^ impact on survival remain incompletely understood.

*DNMT3A* mutations in AML most frequently affect the arginine residue at position 882 (R882), located within the methyltransferase domain ^18^. The R882 variants (e.g. the most common R882H substitution) exert a loss-of-function effect via a dominant-negative mechanism on the wild-type DNMT3A, reducing catalytic activity and impairing CpG methylation ^19-22^. DNMT3A is a key *de novo* DNA methylase active in the body, and serves to inactivate embryonic stem cell programs through promoter methylation ^23^. Concordant with this function, *DNMT3A*^*R882H*^ AMLs display focal hypomethylation specifically at promoter regions ^4,24^ that are linked to hematopoietic stem cell (HSC) self-renewal ^21,25-28^. Prior studies have also attributed *DNMT3A*^*R882H*^ to impaired nucleosome eviction and the attenuation of DNA damage in AML cell lines in response to anthracyclines ^15^. Furthermore, cell lines engineered to ectopically express *DNMT3A*^*R882H*^ exhibit defective DNA repair and replication-fork recovery, leading to accumulation of DNA breaks carried through mitosis and increased *in vitro* sensitivity to cytarabine ^29^. Yet it is unclear if these mechanisms remain clinically relevant in the context of *DNMT3A*^*mut*^*-NPM1*^*c*^*-FLT3*^*ITD*^ AML treated with standard of care combination, anthracycline and cytarabine chemotherapy regimens.

To understand the cellular mechanisms underlying adverse prognosis in *DNMT3A*^mut^*-NPM1*^*C*^*-FLT3*^*ITD*^AML, we generated isogenic matched murine models of *Npm1*^*cA/+*^*-Flt3*^*ITD/+*^ with or without *Dnmt3a*^*R878H/+*^ (mouse equivalent to human R882H). We demonstrate that *Dnmt3a*^*R878H*^ mutation augments Leukaemia Stem Cell (LSC) function and quiescence through epigenetically regulated transcriptional programs, and confer resistance to standard chemotherapy. Strategies that bypass LSC quiescence through cell cycle independent combination regimens, or can overcome quiescence-associated chemotherapy resistance by cell cycle priming may represent a readily tractable approach to improve clinical outcomes in patients with adverse risk AML.

### Material and Methods

#### Generation of *Dnmt3a*^*R878H*^*-Npm1*^*c*^*-Flt3*^*ITD*^ leukemias

Mouse experiments were approved by the QIMR Berghofer Medical Research Institute Animal Ethics Committee (A2109-620). Congenic *Ptprc*^*a/b*^ (CD45.1+/CD45.2+) recipient mice were irradiated with whole-body γirradiation (11 Gy in two doses). Male and Female recipients were transplanted via tail vein injection with 1 × 10^6^ bone marrow (BM) cells from either *Rosa26-Cre*^+/-^*-Dnmt3a*^*R878H/+*^*-Npm1*^*cA/+*^*-Flt3*^*ITD/+*^ (DNF) or *Rosa26-Cre*^+/-^*-Npm1*^*cA/+*^*-Flt3*^*ITD/+*^ (NF) in 200ul Leibovitz’s L-15 medium (Thermo Fisher Scientific, Cat# 11415064).

Following transplantation, recipient mice received Baytril 50 (100 mg/L; Elanco) in drinking water for 2 weeks. Four weeks post-transplantation, Cre-mediated recombination of the floxed *Npm1* allele was induced by administering tamoxifen-containing chow (400mg/kg; Specialty Feeds, Cat#SF09-089) to activate *Npm1*^*cA*^ expression. Recombination was validated by genomic DNA PCR. Engraftment was confirmed by flow cytometry. Upon AML development mice were culled for analysis.

For secondary transplantation experiments, primary DNF and NF AML splenocytes (1 × 10^6^) were transplanted via tail vein injection into sub-lethally irradiated (5.5 Gy) *Ptprc*^*a/b*^ recipients. Sorted populations included MPP3 cells (CD45.2^+^, Lin^-^ Sca1^+^ c-Kit^+^ (LSK), CD48^+^, CD150^-^), MPP2 cells (CD45.2^+^ LSK, CD48^+^, CD150^+^) and GMPs (CD45.2^+^, Lin^-^ c-Kit^+^ (LK), CD16/32^+^, CD34^+^). Donor cell engraftment was assessed for pre-treatment stratification.

#### Chemotherapy regimens

Cytarabine (AraC; 50 mg/kg) and doxorubicin (Dox; 1.5 mg/kg) were administered intravenously for three consecutive days. Alternatively, mice were treated with venetoclax (Ven; 100 mg/kg oral gavage) and/or azacitidine (Aza; 2 mg/kg intraperitoneally) using a shortened 5+2 dosing schedule designed to model the clinical 28-day regimen. Combination pegIFNα– chemotherapy experiments were performed by administering Dox+AraC 48 h following pegIFNα treatment. Full details of drug preparation, dosing schedules, and administration are provided in the Supplementary Methods.

#### Gene expression and epigenetic profiling

Bulk RNA sequencing used the NEBNext® Ultra™ II RNA Library Prep Kit (New England Biolabs) or the SMART-Seq® mRNA LP Kit (Takara Bio). Single-cell gene expression profiling was performed using the Chromium Fixed RNA Profiling Kit (10x Genomics). For epigenetic profiling, chromatin accessibility was assessed by ATAC-seq using the ATAC-Seq Kit (Active Motif), and DNA methylation profiling was performed using the NEBNext® Enzymatic Methyl-seq Kit (New England Biolabs). Detailed library preparation protocols, sequencing parameters, and analysis pipelines are provided in the Supplementary Methods.

#### Statistical Analysis

Statistical analyses were performed using Prism (GraphPad Software, version 10). Normality of data distribution was assessed using the Shapiro–Wilk test. For two-group comparisons, parametric Welch’s t-test or non-parametric Mann–Whitney test was applied. For comparisons involving multiple groups, one-way ANOVA was used, as indicated in the figure legends. Statistical significance is represented as follows: ns, not significant, ^#^p=0.06, *p<0.05, **p<0.01, ***p<0.001, ****p<0.0001.

## Results

### Leukemia Stem Cells (LSCs) are enriched within the multipotent progenitor compartment in *Npm1*^*cA*^-*Flt3*^*ITD*^ and *Dnmt3a*^*R878H/+*^*-Npm1*^*cA*^-*Flt3*^*ITD*^ AML

To isolate the impact of *Dnmt3a*^*R878H*^ mutation when co-occurring with *Npm1*^*cA*^ and *Flt3*^*ITD*^ mutations, we developed isogenic strains harboring *Dnmt3a*^*R878H/+*^*-Npm1*^*cA/+*^*-Flt3*^*ITD/+*^ (DNF) or *Npm1*^*cA/+*^*-Flt3*^*ITD/+*^ mutations with wild-type *Dnmt3a* (NF) (Figure 1A; Supplemental Figure 1A-B). Transplantation of DNF and NF bone marrow (BM) resulted in development of a myelomonocytic AML with short latency in both genotypes (Figure 1B-D). BM composition in both NF and DNF mice demonstrated an increased frequency of monocytes (Mac1^+^) ^14,15^ (Figure 1E). Both DNF and NF showed increased frequency of primitive Lineage-, Sca1+ and c-kit+ (LSK), which almost exclusively comprised the MPP3 (CD48+, CD150-) multipotent progenitor population at the expense of other LSK subgroups, e.g. HSC-SLAM (Figure 1F and G; Supplemental Figure 1C-D), concordant with previous findings for NF AML ^14^. DNF AMLs also showed expansion of Lineage^−^Sca1^−^c-Kit^+^ (LK) progenitors, within the common-myeloid progenitor (CMP) and the granulocyte-macrophage progenitor (GMP) compartment (Supplemental Figure 1C-D).

**Figure 1.**
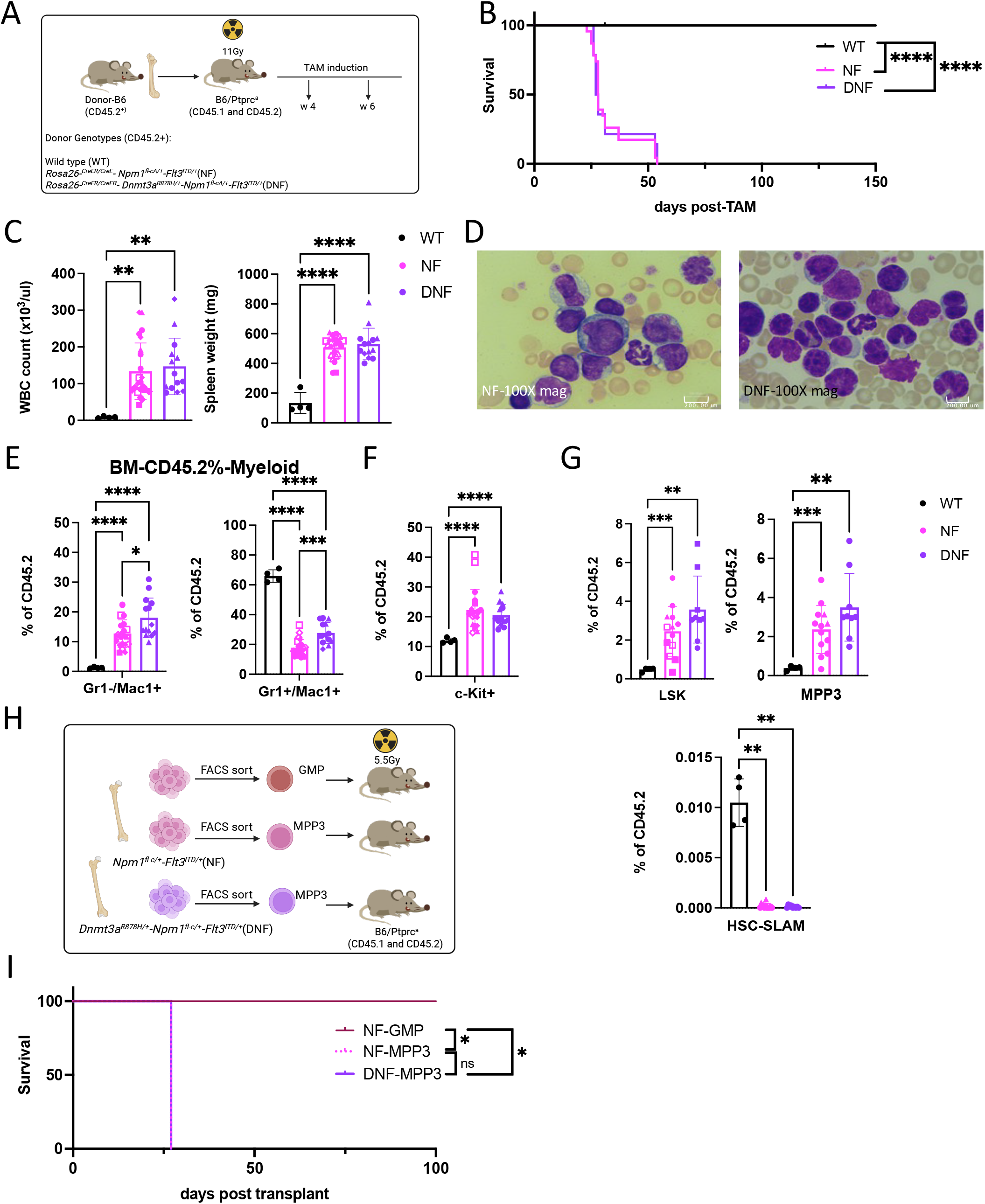
Leukemia stem cells (LSCs) reside within the MPP3 population in *Npm1*^*c*^*-Flt3*^*ITD*^ AMLs with or without *Dnmt3a*^*R878H*^ mutation. (A) Schematic illustration of the experimental strategy used to generate isogenic *Dnmta*^*R878H*^*-Npm1*^*c*^*-Flt3*^*ITD*^ and *Npm1*^*c*^*-Flt3*^*ITD*^ AMLs. (B) Kaplan-Meier survival analysis of mice transplanted with bone marrow (BM) cells from *Rosa26-Cre*^+/-^*-Dnmt3a*^*R878H/+*^*-Npm1*^*cA/+*^*-Flt3*^*ITD/+*^ (DNF; n=14), *Rosa26-Cre*^+/-^*-Npm1*^*cA/+*^*-Flt3*^*ITD/+*^ (NF; n=23) or Wild Type (WT; n=8) donors following tamoxifen (TAM)-induced Cre activation. (C) Peripheral blood (PB) white blood cell (WBC) counts and spleen weights of leukemic DNF and NF mice compared with WT mice (n=4), 6 weeks post-TAM induction without evidence of disease. (D) Representative hematoxylin and eosin (H&E) staining of peripheral blood smears from leukemic DNF and NF mice. Images were acquired using a Nikon Eclipse Ci microscope equipped with XCAMTOP4K camera at 1000× magnification. Scale bar, 200 μm. Flow cytometric analysis of mature lineage compartments in CD45.2^+^ AML BM cells showing the frequency of (E) monocytes (Gr1^-^/Mac1^+^) and granulocytes (Gr1^+^/Mac1^+^), (F) c-Kit^+^. (G) Flow cytometric analysis of hematopoietic stem and progenitor populations (HSPC) demonstrating the frequency of LSK, MPP3, and HSC-SLAM subsets within CD45.2^+^ AML BM cells from DNF and NF mice compared with WT controls (WT vs NF vs DNF, n=4 vs 23 vs 14). (H) Experimental design of secondary transplantation assays using FACS-sorted granulocyte–megakaryocyte progenitors (GMPs) and multipotent progenitor 3 (MPP3) populations isolated from primary DNF and NF AML BM samples. (I) Kaplan–Meier analysis of leukemia development in recipient mice transplanted with NF GMP (n=2), NF MPP3 (n=3), or DNF MPP3 (n=3) cells. Each data point shape represents a biological replicate; identical shapes denote technical replicates derived from the same biological replicate. (B, I) Statistical significance was determined using the log-rank test. (C, E-G) Statistical analyses were performed using ordinary one-way ANOVA. Error bars represent mean ± SD. *p<0.05, **p<0.01, ***p<0.001 and ****p<0.0001.

Secondary transplantation into sub-lethally irradiated recipients confirmed AML transplantation with comparable disease latency (Supplemental Figure 1E). DNF AMLs demonstrated increased frequency of c-Kit^+^ and MPP3 cells, compared with NF (Supplemental Figure 1F), consistent with *Dnmt3a*^*mut*^ causing expansion of the HSC pool over serial transplantation ^21,25^. Additionally, DNF primary AML cells showed enhanced serial replating capacity *in vitro*, suggesting increased self-renewal (Supplemental Figure 1G). To delineate the LSC-enriched population, we isolated BM MPP3 and progenitor (GMP) populations from primary recipients and transplanted these into sub-lethally irradiated recipients (Figure 1H). Both NF and DNF MPP3 populations rapidly induced AML in secondary recipients, however transplantation of more mature GMPs did not, confirming the MPP3 population is highly enriched for functional LSCs (hereafter referred to as LSCs) (Figure 1I).

### *Dnmt3a*^*R878H*^ drives a transcriptional program regulating stem cell function and quiescence in *Npm1*^*c*^*-Flt3*^*ITD*^ LSCs

To distinguish whether the expansion of LSCs in DNF AML reflects an increased LSC frequency or intrinsic differences in self-renewal properties, we performed limiting dilution transplantation assays using MPP3 cells (Supplemental Figure 1H), with no significant differences between LSC frequency in NF and DNF (Supplemental Figure 1I). Next, we assessed whether *Dnmt3a*^*R878H*^ alters gene expression of LSCs in NF AML by performing RNA-seq on purified LSC-enriched MPP3 cells from NF and DNF leukemias (Figure 2A; Supplemental Table 2). Gene Set Enrichment Analysis (GSEA) showed significant enrichment of transcriptional signatures associated with LSC phenotype (NES=2.3, FDR<0.0001) and quiescent LSCs ^30^ (NES=1.9, FDR=0.001) in DNF compared to NF LSCs (Figure 2B and Supplemental Figure 2A; Supplemental Table 3). Conversely, *Dnmt3a*^*R878H*^ LSCs exhibited decreased expression of genes associated with differentiation (Supplemental Figure 2B) and MYC targets (Supplemental Figure 2C), supportive of increased stemness and reduced proliferation. These data were confirmed with flow cytometry cell cycle analysis showing increased frequency of quiescent G0 cells (Ki-67 negative population) in DNF compared to NF LSCs (Figure 2C).

**Figure 2.**
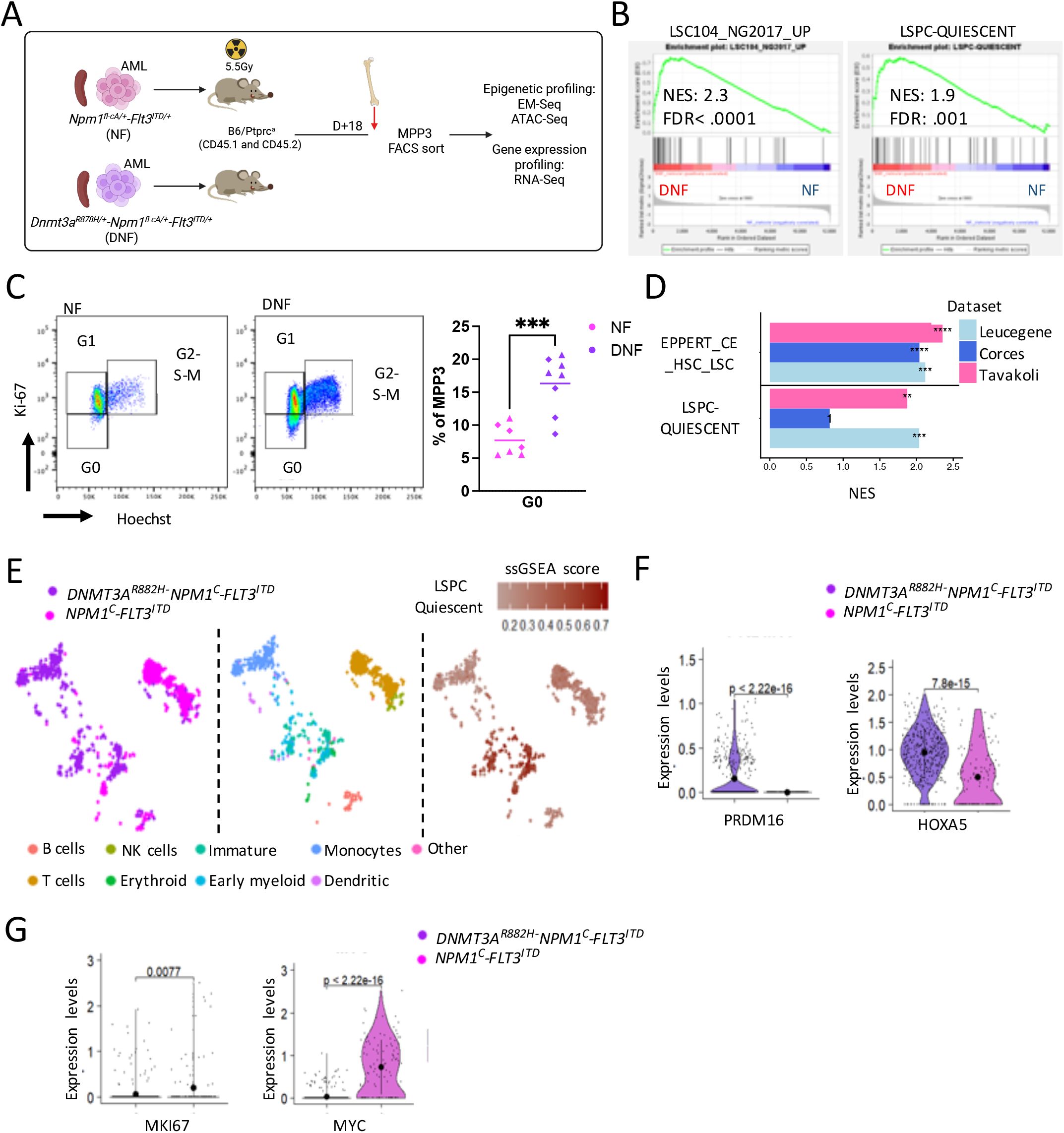
*DNMT3A* mutation induces an enhanced quiescence signature within the LSC population that is conserved in human *NPM1*^*c*^*-FLT3*^*ITD*^ AML. (A) Experimental design for isolation of LSC-enriched MPP3 cells from DNF and NF AML (<20% AML infiltration in BM) followed by bulk RNA-seq, ATAC-seq and EM-seq analyses. (B) Gene set enrichment analysis (GSEA) plots showing normalized enrichment score (NES) and false discovery rate (FDR) for LSC and quiescence-associated gene signatures in DNF (n=4) versus NF (n=4) LSCs. (C) Representative flow cytometry plots of DNF and NF LSC-enriched MPP3 cells stained for the proliferation marker Ki-67 and Hoechst (DNA content) to assess cell cycle status. Quantification of the frequency of G_0_ (quiescent) cells within the LSC-enriched MPP3 population in DNF (n=8) versus NF (n=7). Each symbol represents a biological replicate; identical symbols denote technical replicates derived from the same biological sample. Statistical analyses were performed using unpaired Welch’s t-test. (D) Bar graph showing GSEA NES scores for stemness and quiescence signatures in *DNMT3A*^*R882H*^*-NPM1*^*c*^*-FLT3*^*ITD*^ compared with *NPM1*^*c*^*-FLT3*^*ITD*^ AML across independent diagnostic patient cohorts. Analysis includes Leucegene datasets with genomic annotations from Garg et al. (2019) ^31^ and Corces-Zimmerman et al. (2014) ^32^, comparing *DNMT3A*^*R882H/C*^*-NPM1*^*c*^*-FLT3*^*ITD*^ (n=5) versus *NPM1*^*c*^*-FLT3*^*ITD*^ (n=4), and *DNMT3A*^*mut*^*-NPM1*^*c*^*-FLT3*^*ITD*^ (n=2) versus *NPM1*^*c*^*-FLT3*^*ITD*^ (n=2) diagnostic samples, respectively. Human gene signatures were compared with DNF versus NF murine LSC gene signatures generated in this study. FDR significance levels are indicated as asterix **<0.01, ***<0.001, ****<0.0001. (E) Uniform Manifold Approximation and Projection (UMAP) of diagnostic single-cell RNA-Seq data from *DNMT3A*^*R882H*^*-NPM1*^*c*^*-FLT3*^*ITD*^ and *NPM1*^*c*^*-FLT3*^*ITD*^ AML patient samples ^33^ color-coded (left to right) by genotype, cluster annotation, and ssGSEA derived quiescence signature scores. Violine plots comparing the normalized gene expression levels of (F) *HOXA5* and *PRDM16*, and (D) *MYC* and *MKI-67* in immature AML cells from *DNMT3A*^*R882H*^*-NPM1*^*c*^*-FLT3*^*ITD*^ versus *NPM1*^*c*^*-FLT3*^*ITD*^ AML patients. (F-G) Statistical significance was determined by pairwise, two-sided Welch’s-t-test-with black dot and error bars indicate mean ± SD, respectively. ***P<0.001.

To validate these findings in human *DNMT3A*^*mut*^*-NPM1C-FLT3*^*ITD*^ AML, we used gene expression profiles of diagnostic AML patient samples from the publicly available datasets ^31,32^. *DNMT3A* mutations were associated with increased expression of LSC- and quiescence-associated gene signatures in *NPM1*^*C*^*-FLT3*^*ITD*^ human AML (Figure 2D). In human single-cell RNA-sequencing datasets^33^, *DNMT3A*^*R882H*^*-NPM1*^*c*^*-FLT3*^*ITD*^ CD34^+^ cells displayed a higher frequency of cells occupying immature transcriptional compartments compared with *NPM1*^*c*^*-FLT3*^*ITD*^ controls (Figure 2E). Within this immature population, *DNMT3A*-mutant cells exhibited increased quiescence signatures, accompanied by elevated expression of key stemness regulators including *PRDM16*, a critical mediator of long-term HSCs maintenance and quiescence ^34,35^, and HOX genes such as *HOXA5*, that are associated with HSC self-renewal regulation and LSC identity ^36,37^ (Figure 2F). Consistent with a quiescent state, these cells also exhibited reduced expression of proliferation markers such as *MKI67* and *MYC* (Figure 2G). Supporting this observation, independent studies have reported *PRDM16* overexpression in *DNMT3A*-mutant *NPM1*^*c*^ AML, where elevated PRDM16 expression is associated with inferior overall survival in *NPM1*^*c*^*-FLT3*^*ITD*^ AML ^38,39^. Together, these data validate our murine observations and identify a *DNMT3A*-driven quiescent LSC program in human AML.

### *Dnmt3a*^*R878H*^ mutation causes epigenetic dysregulation leading to aberrant gene expression and increased stem cell quiescence

To determine whether the transcriptional changes in DNF LSCs are a direct consequence of loss of *DNMT3A* methyltransferase function, we performed Enzymatic Methyl-seq (EM-seq) on the LSC-enriched MPP3 population. Hierarchical clustering on the basis of CpG methylation separated *Dnmt3a*^*R878H*^ mutated NF LSCs from *Dnmt3a*^*WT*^ (Supplemental Figure 3A), indicating a measurable difference in DNA methylation across genotypes. Out of 14,181,562 CpG sites analyzed, 1,740,049 (12.2%) were differentially methylated in DNF compared to NF LSCs. These differentially methylated regions (DMRs) were predominantly associated with loss of methylation (1,352,649; 74.7%), whereas a small portion showed gain of methylation (457,862, 25.3%) (Figure 3A; Supplemental Table 4). Although the DNA hypomethylation was noticeable across all regions of the genome, difference in methylation was greatest at gene promoters, CpG islands and shores (Figure 3B, Supplemental Figure 2E). However, at the genome wide level, changes in promoter methylation were not independently predictive of gene expression (Supplemental Figure 2F).

**Figure 3.**
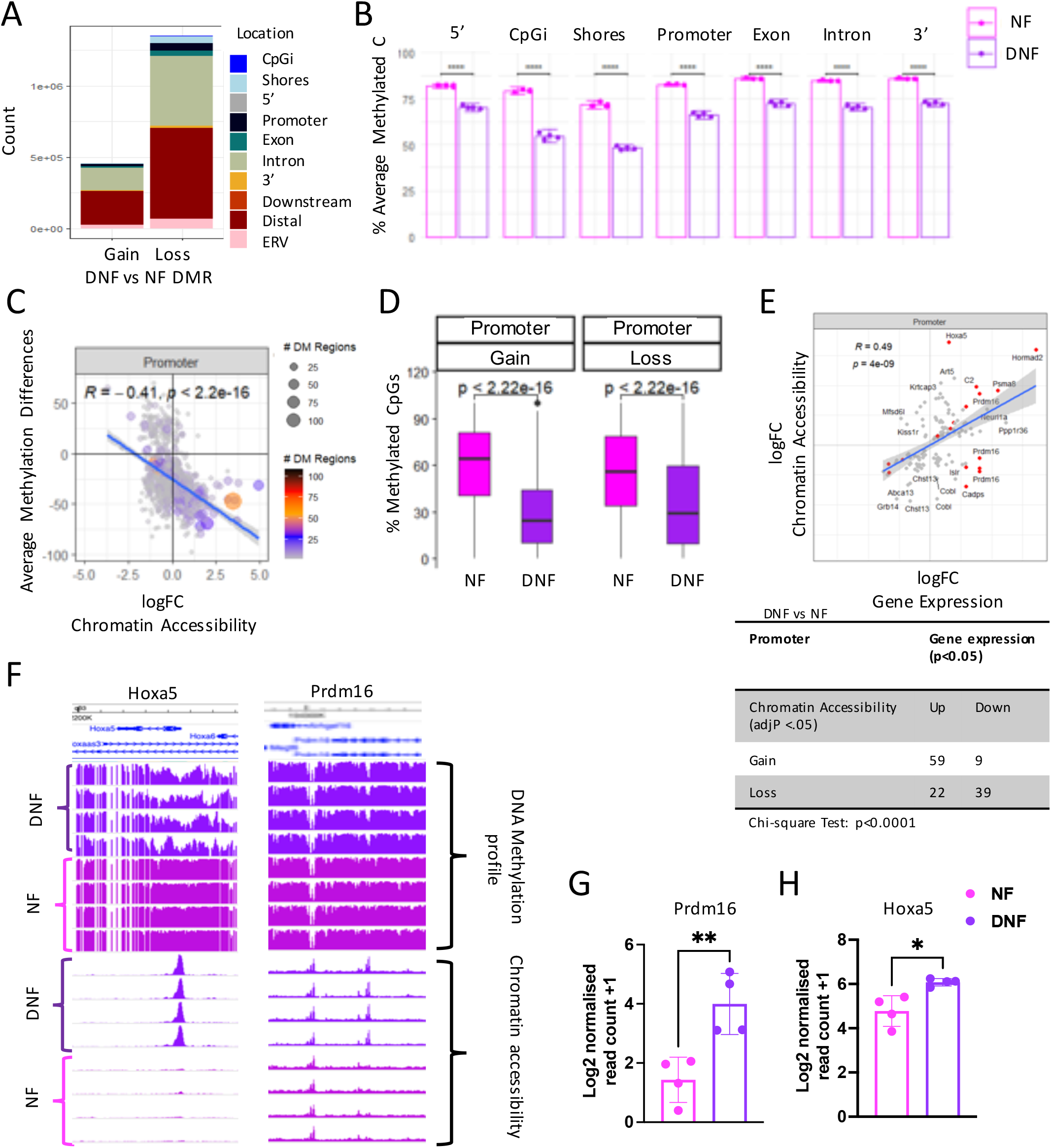
*Dnmt3a*^*R878H*^ mutation causes reduced DNA methylation and increased chromatin accessibility at promoters of genes associated with quiescence in LSCs. (A) Bar graph displaying the number of gained and lost differentially methylated regions (DMR; FDR <0.05) in DNF (n=4) versus NF (n=4) LSCs across indicated genomic regions. (B) Average percentage of EM-seq derived DNA methylation in DNF versus NF LSCs across genomic features. (C) Scatterplot of differential promoter-associated ATAC-seq peaks (FDR<0.05) in DNF versus NF LSCs, showing log fold change (logFC) in chromatin accessibility versus corresponding differences in average DNA methylation at overlapping DMRs (FDR<0.05). Color and size indicate the number of differently methylated CpGs (FDR<0.05) under the ATAC peak. (D) Boxplots displaying the percentage of methylated CpGs at differentially ATAC gained or lost promoters (FDR<0.05) in DNF compared with NF LSCs. (E) Scatter plot showing gene expression changes (logFC) compared to differences in chromatin accessibility (ATAC-seq read coverage; logFC) in DNF versus NF LSCs at promoter region of significantly differentially expressed genes (FDR<0.05). Red dots indicate genes with significantly differential ATAC peaks (FDR<0.05). Table summarizes the number of differentially up- or down-regulated genes (p<0.05) with significant differences in ATAC peak read coverage (FDR <0.05) and chi-square test of independence p-value. (F) The percentage of cytosine methylation (range 0-100%) and normalized ATAC-seq read coverage at the *Prdm16* and *Hoxa5* loci in DNF and NF LSCs, visualized in the WashU Epigenome browser. Bar heights represent the mean. Bulk RNA-seq derived normalized expression of (G) *Prdm16* and (H) *Hoxa5* in DNF compared to NF LSCs. Data includes four independent replicate per genotype. (B, D, G-H) Statistical analysis was performed by two-sided, pairwise Welch’s Student T-test. (C, E) Display coefficients (R) and p-values of a Pearson’s correlation test. Error bars represent mean ± SD.*P<0.05, **p<0.01 and ****p<0.0001.

To further delineate the epigenetic profile induced by *Dnmt3a*^*R878H*^ mutation in NF AML, we profiled chromatin accessibility of DNF and NF MPP3s using ATAC-seq. Consistent with the DNA methylation differences, principal component analysis (PCA) of chromatin accessibility across all genomic regions, and specifically at promoters, revealed that DNF LSCs cluster separately from NF LSCs (Supplemental Figure 2G). Changes in chromatin accessibility correlated negatively with changes in DNA methylation across all genomic regions, including promoters (R=-0.33, p<2.2e-16; Figure 3C and Supplemental Figure 2H). Gain of promoter accessibility was widely associated with a loss of DNA methylation in DNF compared to NF LSCs (Figure 3D). Paradoxically, a subset of promoters in DNF LSCs showed concurrent loss of accessibility and decreased methylation (Figure 3D), which may reflect a lag phase due to impaired or delayed histone eviction ^15^. Importantly, genes that had increased expression in DNF LSCs were associated with increased promoter chromatin accessibility, while downregulated genes were predominantly linked to reduced promoter accessibility (Chi-square p<0.0001; Figure 3E). Furthermore, hypomethylated promoter regions in DNF compared to NF LSCs were significantly enriched for genes associated with HSC and stemness, while genes contained in hypermethylated promoters were involved in myeloid progenitor differentiation and cell division (Supplemental Figure 2I). Specific examples of genes that demonstrated increased expression in DNF, together with increased chromatin accessibility and hypomethylation at promoter regions include the quiescence regulator, *Prdm16* ^34,35^, and *Hoxa5* ^36,37^ (Figure 3F-H), also increased in human DNF AML (Figure 2F).

These data demonstrate that *Dnmt3a*^*R878H*^ diminishes DNA methylation, leading to increased chromatin accessibility at promoter regions that regulate a LSC-enriched, quiescence gene expression program.

### *Dnmt3a*^*R878H*^-mutated *Npm1*^*c*^*-Flt3*^*ITD*^ LSCs are resistant to cytotoxic chemotherapy

We next sought to compare the response to combined chemotherapy in DNF versus NF AMLs. Recipient mice were treated with 3 doses of doxorubicin and cytarabine (Dox+AraC) combination (Figure 4A), modelling the chemotherapy combination that forms the backbone of clinically used standard chemotherapy regimens for AML patients ^40,41^. NF mice showed effective clearance of AML cells from peripheral blood, spleen and BM following Dox+AraC treatment, whereas DNF AML were resistant to chemotherapy in bulk AML (Figure 4B-D) and within phenotypically defined LSC compartments (Figure 4E).

**Figure 4.**
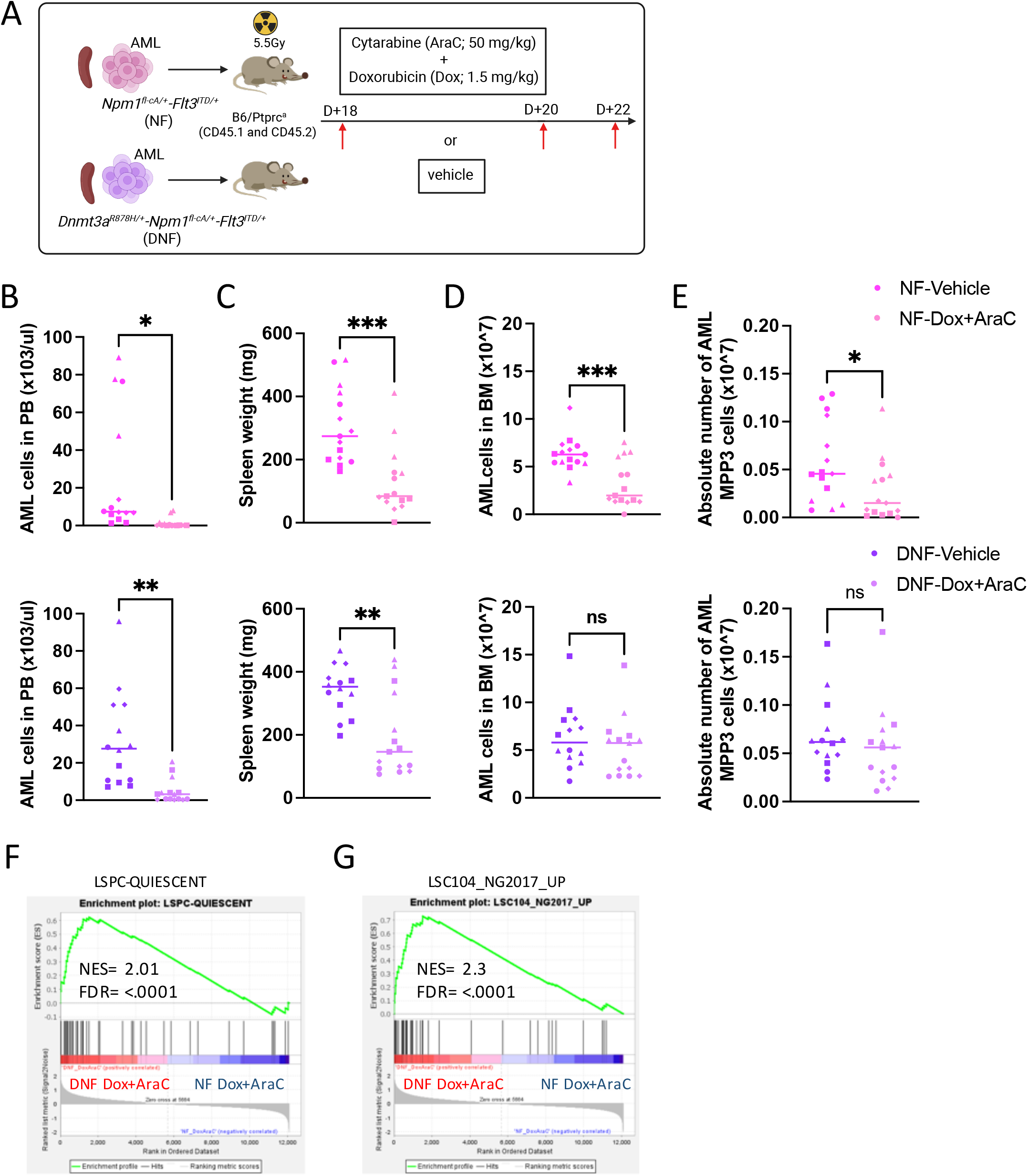
Standard chemotherapy fails to deplete the *Dnmt3a*^*R878H*^-mutated LSCs. (A) Experimental schema for treatment of DNF and NF mice with standard chemotherapy (three doses of doxorubicin plus cytarabine; Dox+AraC) or vehicle control. Quantification of (B) Circulating PB AML blasts, (C) Spleen weight, (D) BM AML burden and (E) LSC-enriched MPP3s in NF and DNF AMLs following vehicle (NF, n=15; DNF, n=14) or Dox+AraC (NF, n=15; DNF, n=15) treatment. Each data point represents an independent biological replicate with median indicated by the horizontal line. Identical symbols denote technical replicates derived from the same biological sample pooled across four independent experiments. GSEA with NES and FDR of bulk RNA-seq from DNF versus NF LSCs after Dox+AraC treatment showing (F) quiescence- and (G) LSC-associated transcriptional signatures (n=4 biological replicates per genotype and treatment). (B-E) Statistical significance was determined using an unpaired Mann-Whitney test. *p<0.05, **p<0.01 and ***p<0.001.

To determine whether this chemoresistance reflected impaired DNA damage response of *DNMT3A*-mutant cells ^15^, we assessed double strand DNA breaks following Dox+AraC treatment. *Ex vivo*, treatment induced comparable γH2AX phosphorylation in DNF and NF AML cells (Supplemental Figure3 A-B). *In vivo*, bulk DNF AML showed reduced DNA damage compared with NF following treatment (Supplemental Figure 3C), however, γH2AX levels within the LSC-containing LSK compartment were similar between genotypes at baseline (T0) and 48 hours post-treatment (T48) (Supplemental Figure 3D). These results were supported by transcriptional profiling of LSCs 48 hours post chemotherapy which demonstrated comparable induction of p53 and DNA repair pathways in NF and DNF LSCs (Supplemental Figure 3E). Importantly, DNF LSCs retained higher quiescence and stemness gene signature after chemotherapy when compared to NF LSCs (Figure 4F-G; Supplemental Figure 3F). These data indicate that resistance to combination chemotherapy in DNF LSCs is not primarily driven by defective DNA damage sensing, but rather reflects persistence of protective quiescent stem cell state.

### Chemotherapy resistance in *Dnmt3a*^*R878H*^ AML resides in a transcriptionally distinct quiescent LSC population

To dissect the LSC heterogeneity underlying chemotherapy resistance and identify the resistant LSC subpopulation in DNF AML, we performed single cell gene expression profiling on DNF and NF LSCs treated with, or without chemotherapy (Figure 5A). Uniform Manifold Approximation and Projection (UMAP) analysis revealed heterogeneity within the LSC compartment of both vehicle-treated AMLs (Figure 5B). Transcriptionally defined LSC clusters correspond to hierarchical hematopoietic states spanning primitive HSC-like populations, multipotent progenitors, lineage-biased progenitors and clusters showing activation of transcriptional features associated with mature cell differentiation (Supplemental Figure 4A-B) ^30,42-44^. In comparison to WT LSK (Supplemental Figure 3C), DNF and NF LSC populations demonstrate the over-representation of MPP3-like clusters, supporting the surface marker expression analysis (Figure 1H and Supplemental Figure 1D).

**Figure 5.**
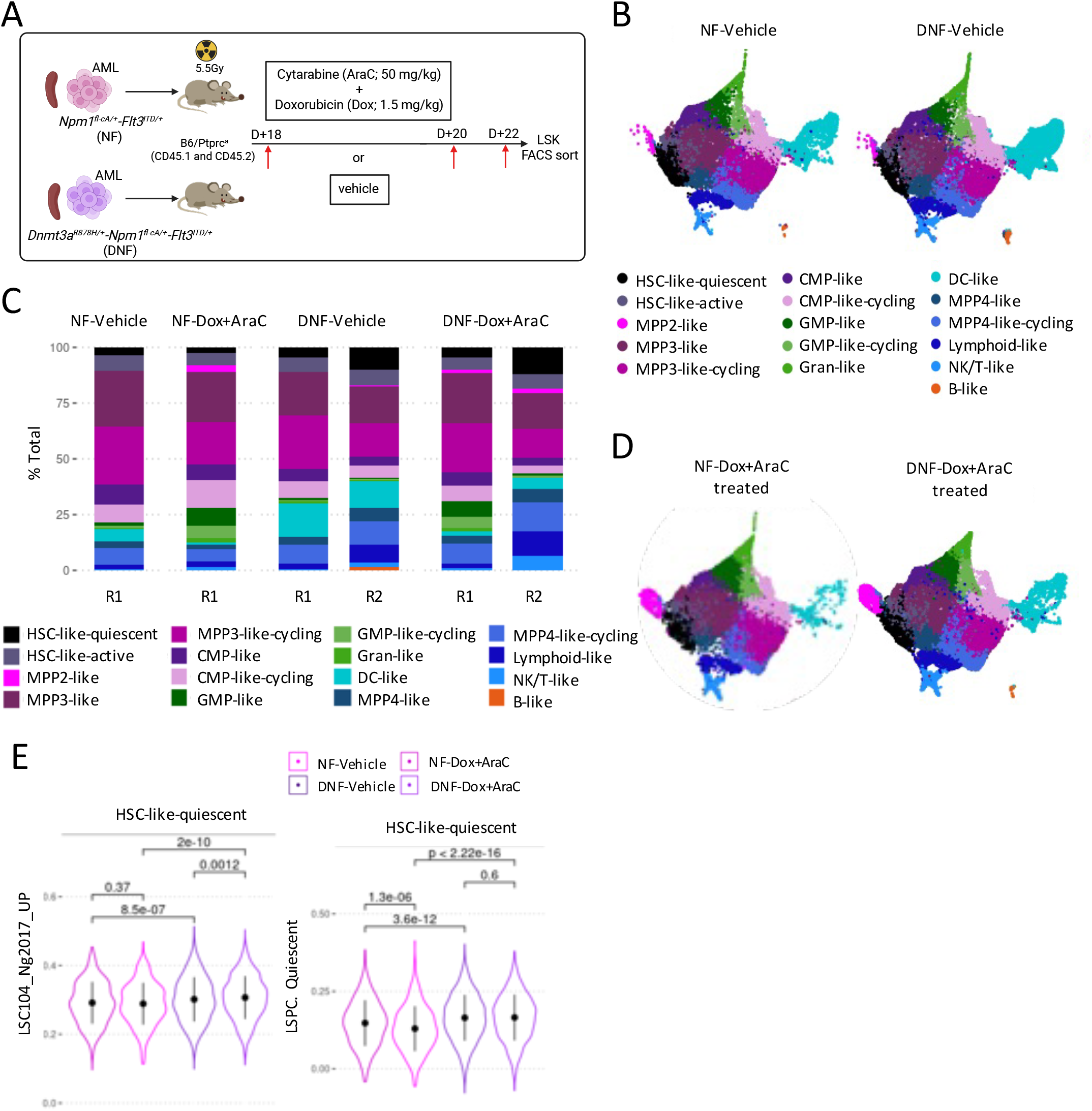
*Dnmt3a*^*R878H*^-driven chemoresistance is concentrated within the primitive HSC-like LSC population in *Npm1*^*c*^*-Flt3*^*ITD*^ AML. (A) An experimental schema for isolation of LSC-containing LSK cells from vehicle- or Dox+Arac-treated DNF and NF AMLs for single-cell gene expression profiling. (B) UMAP of single-cell RNA-seq data from NF and DNF AML treated with vehicle control (NF, n=1; DNF, n=2 pooled independent biological replicates). Each point represents a single cell, colored by transcriptionally defined clusters. (C) Bar graph showing the proportion of transcriptionally defined clusters in NF and DNF LSC-containing LSKs treated with vehicle or Dox+AraC with colors as defined in B. R1 and R2 denote independent biological leukaemias processed in separate experimental runs. (D) UMAP of single-cell RNA-seq data from DNF and NF AMLs treated with Dox+AraC. Violin plots of single cell GSEA derived (E) LSC gene signature, and quiescence scores within the HSC-like quiescent cluster in NF and DNF LSC-containing LSKs under vehicle or Dox+AraC treatment. P-values determined using two-sided, unpaired Welch’s T-test. Dots and error bars represent as mean ± SD, respectively.

Consistent with the bulk RNA sequencing analysis (Figure 2B and Figure 4F-G), higher expression of LSC and quiescent gene signatures were found in DNF compared with NF, both in the absence and presence of chemotherapy (Supplemental Figure 4D-E). A primitive HSC-like population that showed distinct enrichment of quiescent stem cell programs was termed HSC-like quiescent (Supplemental Figure 4A-B). After chemotherapy treatment, the HSC-like quiescent cluster was selectively preserved in DNF AML (vehicle vs Dox+AraC-treated; Figure 5C-D). Although chemotherapy reduced LSC and quiescence programs globally (Supplemental Figure 4D-E), these signatures were enriched or maintained within the DNF HSC-like quiescent LSC cluster, but not in NF LSCs (Figure 5E). Cell-cycle analysis further confirmed that this population remained predominantly in G0-G1 following chemotherapy treatment (Supplemental Figure 4F). Together, these findings identify a chemotherapy resistant, quiescent LSC population in DNF AML that acts as a reservoir for AML relapse.

### *Dnmt3a*^*R878H*^ LSCs respond to cycle independent venetoclax and azacitidine combination therapy

The BCL2 inhibitor, venetoclax, is used in combination with azacitidine for patients with AML who are unfit for intensive chemotherapy (Ven+Aza) ^45,46^. As our data support the resistance of DNF LSCs to chemotherapy through enhanced quiescence, and as venetoclax and azacitidine act largely independent of cell-cycle ^40,47,48^, we next evaluated the response of *Dnmt3a*^*R878H*^-mutated NF AML to Ven+Aza.

We developed and validated an *in vivo* Ven+Aza regimen ^46^ for a total of 7 days (Figure 6A). Ven+Aza resulted in depletion of circulating blasts and substantial reductions in AML burden within spleen and BM in both NF and DNF mice (Figure 6B-D). Notably, Aza-treatment alone and/or in combination induced a granulocytic differentiation phenotype in DNF and NF BM (Supplemental Figure 5A-B), consistent with previous reports ^49-52^, and this was accompanied by a significant reduction in myeloid blasts (Supplemental Figure 5C-D). Treatment with either single-agent Ven and Aza was sufficient to reduce the circulating AML cells in peripheral blood (Supplemental Figure 5E); however, Ven alone had no effect on AML burden in the BM (Supplemental Figure 5F-G).

**Figure 6.**
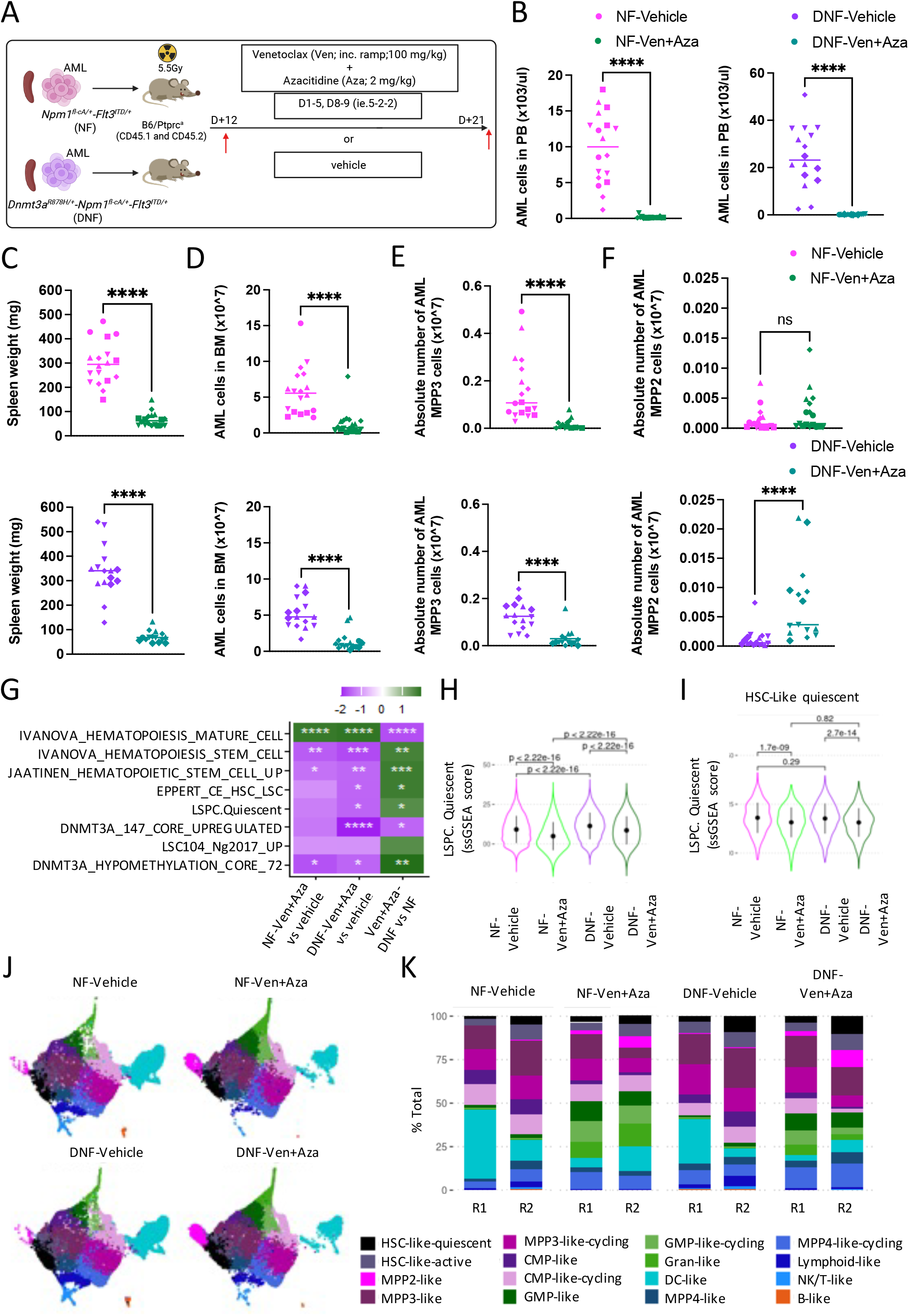
Venetoclax-based combination therapies with azacitidine is effective in depleting *Dnmt3a*^*R878H*^ LSCs in *Npm1*^*c*^*-Flt3*^*ITD*^ AML. (A) Experimental setup for a 7-day randomized treatment regimen with venetoclax plus azacitidine (Ven+Aza), venetoclax alone (Ven), azacitidine alone (Aza), or vehicle control. Disease burden in NF and DNF AML mice treated with Ven+Aza (NF, n=18; DNF, n=16) or vehicle control (NF, n=18; DNF, n=15) measured by (B) PB leukocytes, (C) spleen weight, (D) BM AML blasts, and (E-F) LSC-enriched MPP3 and MPP2 within the c-Kit^+^ positive primitive population. Each data point represents an independent biological replicate with median indicated by the horizontal line. Identical symbols indicate technical replicates derived from the same biological sample, pooled across five independent experiments. (G) Heat map GSEA NES of indicated gene signatures comparing gene expression from bulk RNA-seq in Ven+Aza-treated vs vehicle-treated NF LSCs, Ven+Aza-treated vs vehicle-treated DNF LSCs and Ven+Aza-treated DNF vs NF LSCs (left to right; NF, n=2, DNF, n=2 independent biological replicates per treatment). Green denotes positive enrichment and purple negative enrichment. Statistical significance from GSEA is indicated by asterisks, *FDR<0.05, **FDR<0.01, ***FDR<0.001 and ****FDR<0.0001. Violin plots of single-cell gene expression GSEA of the LSPC quiescent signature scores (H) across all data, or (I) within the HSC-like quiescent clusters in NF and DNF LSC-containing LSKs following Ven+Aza or vehicle treatment. Dot and error bars represent as mean ± SD, respectively. (J) UMAPs of NF and DNF LSC-containing LSKs after Ven+Aza or vehicle treatment. (NF, n=2, DNF, n=4 independent biological replicates per treatment pooled into 2 runs). (K) Proportion of NF and DNF LSKs across different transcriptional clusters upon Ven+Aza or vehicle treatment. R1 and R2 denote independent biological leukaemias processed in separate experimental runs. Statistical significance was determined using an unpaired Mann Whitney test (B-F) or a two-sided, unpaired Welch T-test (H-I). ***p<0.001.

In contrast to DNF LSC resistance to Dox+Ara chemotherapy, NF and DNF showed comparable reductions in LSC frequency following Ven+Aza exposure, indicating that DNF-associated resistance is partially overcome by Ven+Aza combinations (Figure 6E). Reduction of the LSC-enriched MPP3 compartment was driven primarily by Ven, particularly in DNF mice (Supplemental Figure 5H), consistent with high expression of BCL2 in primitive progenitor populations ^53-55^. In contrast, Aza alone had predominant effects on bulk AML burden and myeloid blasts in the BM (Supplemental Figure 5F-H). Providing caution to this approach, we observed an increased frequency of the more primitive MPP2 population in DNF mice following Ven+Aza, not seen in NF, although this population was present at lower numbers than the dominant MPP3 population (Figure 6F and Supplemental Figure 5I), suggesting transcriptional plasticity and reversion to a de-differentiated state. Sorted DNF MPP2 cells were capable of engraftment and leukemia initiation (Supplemental Figure 5J) and therefore represent a Ven+Aza resistance mechanism unique to DNF AML.

Bulk RNA-seq analyses performed following Ven+Aza treatment in both NF and DNF LSCs showed strong negative enrichment for stem cell signatures compared to vehicle-treated controls, indicating effective suppression of LSC programs. Furthermore, Ven+Aza treatment was also associated with decreased expression of quiescence-associated genes, particularly in DNF LSCs and attenuation of transcriptional programs previously linked to *DNMT3A* loss ^22^. Despite this, relative to NF, DNF LSCs retained greater enrichment of stem cell and quiescence programs following treatment (Figure 6G). Single cell analysis of the LSK-containing LSC populations further demonstrated that the Ven+Aza treatment reduced the expression of gene sets associated with quiescence, including within the HSC-like quiescent subset (Figure 6H-J). However, this therapy was unable to reduce the frequency of HSC-like quiescent LSCs (Figure 6J-K), suggesting that this combination alone is insufficient to completely overcome the negative prognostic effect of DNF in LSCs.

### Enforced cell-cycle entry sensitizes *Dnmt3a*^*R878H*^ LSCs to chemotherapy

Pegylated interferon-α (pegIFNα) can induce cycling in normal quiescent hematopoietic and *Jak2*^*V617F*^-expressing stem cells in myeloid malignancy ^56,57^. Thus, we next examined whether pegIFNα synergizes with Dox+Ara to induce cell-cycle entry and enhance chemotherapy response in DNF LSCs (Figure 7A). A single dose of pegIFNα had some effect on reducing DNF AML burden and decreased the number of Mac1+ and c-kit+ AML cells in the BM (Supplemental Figure 6A-G), whereas combined pegIFNα and Dox+AraC treatment resulted in a marked reduction of leukemic burden in the spleen (Figure 7B-C), clearance of AML cells from the BM (Figure 7D), depletion of mature monocytic myeloid blasts (Supplemental Figure 6H-I), primitive c-Kit^+^ AML cells (Supplemental Figure 6J-L), and, critically, the LSC-enriched MPP3 population (Figure 7E). An additional clearance of MPP2 cells was also observed (Supplemental Figure 6M). Although pegIFNα alone did not significantly reduce quiescent cells within the LSC compartment (Supplemental Figure 6F), combined pegIFNα and Dox+AraC resulted in marked reduction in the quiescent fraction of DNF MPP3s compared to chemotherapy alone (Figure 7F).

**Figure 7.**
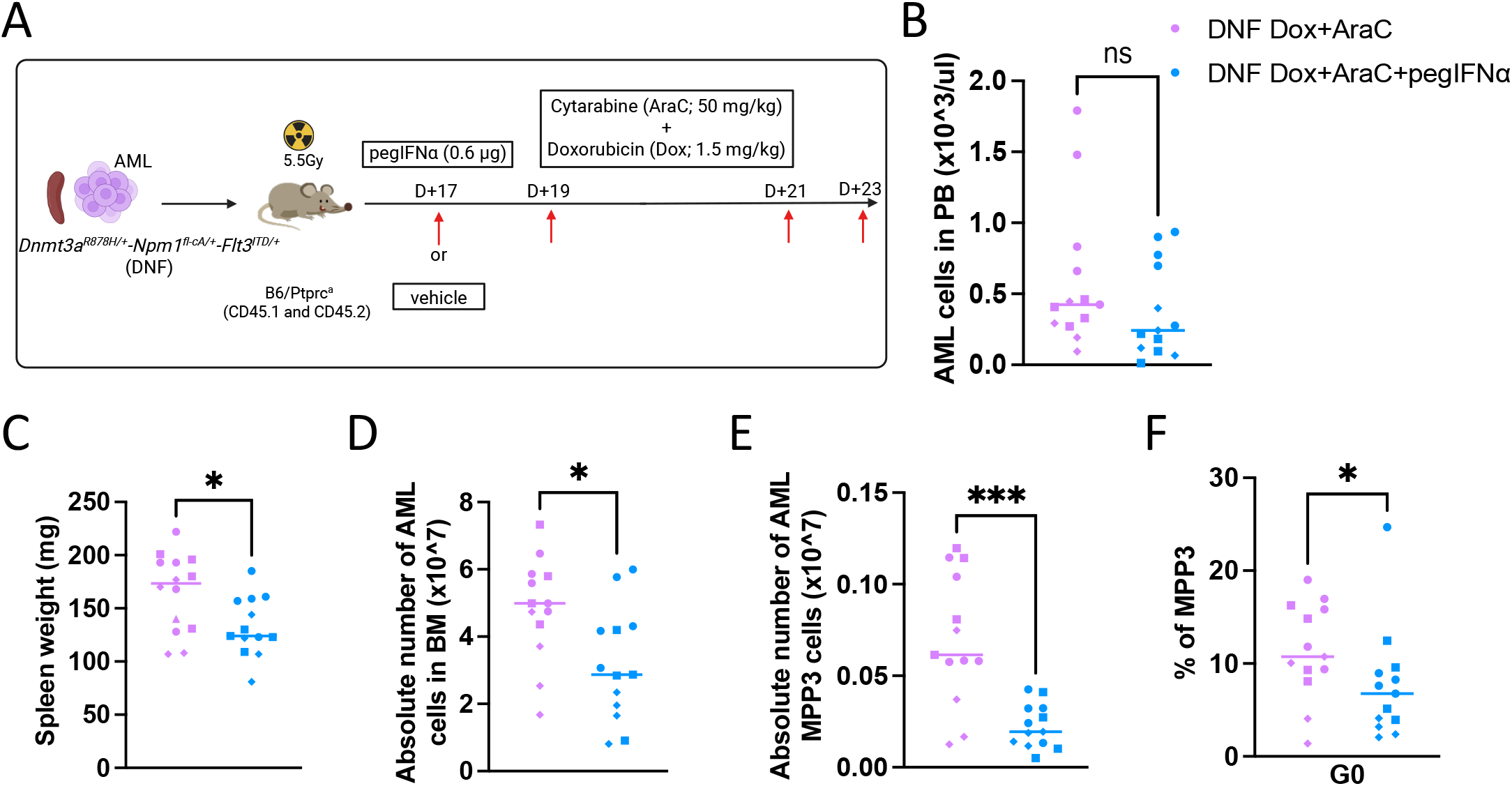
PegIFNα sensitizes *Dnmt3a*^*R878H*^-mutated LSCs to chemotherapy by promoting cell cycle entry. (A) Experimental schema outlining the pegIFNα plus Dox+AraC treatment regimen. DNF mice received a single dose of pegIFNα or vehicle, followed 48 hours later by three doses of Dox+AraC as previously described. AML burden was assessed by (B) number of circulating AML blasts in PB, (C) spleen weight, (D) number of BM AML blasts, and (E) LSC-enriched MPP3 cells in DNF mice pre-treated with pegIFNα prior to chemotherapy. (F) Frequency of quiescent LSC-enriched MPP3s in DNF AML following Dox+AraC + pegIFNα compared with Dox+AraC alone. Each data point represents an independent biological replicate with median indicated by the horizontal line. Identical symbols indicate technical replicates derived from the same biological sample pooled across three independent experiments. Statistical significance was determined using unpaired Mann Whitney test. *p<0.05 and ***p<0.001.

Together, these data validate the concept of LSC priming with pegIFNα prior to chemotherapy. Moreover, these data demonstrate that enforced cell-cycle entry sensitizes DNF LSCs to chemotherapy and identify pegIFNα as a novel strategy to overcome *DNMT3A*-associated treatment resistance in the presence of *NPM1* and *FLT3*^*ITD*^.

## Discussion

Concurrent *DNMT3A, NPM1*, and *FLT3* mutations define a high-risk AML subtype characterized by frequent relapse and inferior survival following chemotherapy ^5,7,8,58^. Despite this recognized clinical paradigm, the mechanisms by which *DNMT3A* mutations drive therapeutic resistance have remained unclear.

Our isogenic models allow isolation of the specific effect of *Dnmt3a* mutation in *Npm1*^*c*^*-Flt3*^*ITD*^ AML. Phenotypically, NF and DNF mice recapitulated the human disease with myelomonocytic morphology and hyperleukocytosis, and displayed comparable disease latency, in contrast to previously published models of inducible *Mx1-Cre-Dnmt3a*^*R878H*^ in which DNF mutations were associated with accelerated disease onset ^15^. Consistent with prior studies ^21,25,59^, the impact of *Dnmt3a*^*R878H*^ mutation in our model was most pronounced within the dominant primitive MPP3 compartment, which functionally harbored the highest leukemia propagating activity and hence, was defined as the LSC compartment.

At the molecular level, integrated transcriptomic and epigenetic analyses demonstrated that *Dnmt3a*^*R878H*^ induces widespread remodeling of transcriptional and epigenetic landscapes in NF LSCs. This included hypomethylation and increased chromatin accessibility at promoter CpG regions associated with stemness and quiescence programs, resulting in sustained expression of LSC- and quiescence-related genes. These findings are supported by recent evidence demonstrating that correction of *DNMT3A*^*R882H*^ mutations in primary *DNMT3A*^*R882H*^-*NPM1*^*C*^*-FLT3*^*ITD*^ AML samples, restores methylation at LSC-associated loci, reduced LSC gene signature expression and decreased functional LSC frequency ^60^. Together, these findings support a model in which *DNMT3A* mutation stabilizes an LSC-enriched, quiescent transcriptional state through epigenetic reprogramming in *NPM1*^*C*^*-FLT3*^*ITD*^ AML.

Prior studies have reported divergent effects of *DNMT3A*^*R882H*^ on chemotherapy response, including attenuated anthracycline-induced DNA damage attributed to impaired nucleosome remodeling and increased cytarabine sensitivity linked to defective replication fork recovery ^15,29^. While these mechanisms implicate altered DNA damage processing in specific experimental contexts, particularly bulk cycling AML cells and cell line models, we did not observe differential DNA damage signaling in DNF LSCs. Instead, we identify maintenance of a quiescent LSC compartment as a central mechanism underlying *Dnmt3a*^*R878H*^-driven chemotherapy resistance in *Npm1*^*C*^*-Flt3*^*ITD*^ AML.

Quiescence has long been associated with chemotherapy resistance ^57,61-63^. Consistent with this, we show that standard Dox+AraC chemotherapy fails to effectively deplete LSCs in *Dnmt3a-*mutated NF AML. While transcriptomic analyses revealed global enrichment of quiescence programs in DNF compared with NF LSCs, single-cell profiling demonstrated that the depth and persistence of quiescence are greatest within a primitive HSC-like LSC population. These analyses reveal that *Dnmt3a*^*R878H*^ reshapes LSC heterogeneity and localizes these epigenetically reinforced quiescent programs to a discrete HSC-like LSC subpopulation, resulting in compartmentalized chemotherapy resistance. The persistence of this quiescent LSC compartment may also provide a mechanistic explanation for the higher rates of measurable residual disease (MRD) frequently observed in *DNMT3A*-mutant *NPM1*^*c*^ AML following chemotherapy, despite equivalent rates of complete morphological remission ^12,17^. Patients with *DNMT3A*^*mut*^*-NPM1*^*c*^*-FLT3*^*ITD*^ AML often remain MRD-positive through induction and consolidation chemotherapy ^12^, suggesting that deeply quiescent LSCs can evade treatment and persist as residual disease, ultimately contributing to relapse.

Recent studies have predicted enhanced sensitivity of *DNMT3A*-mutated AML to venetoclax-based therapies with azacitidine, potentially reflecting increased BCL-2 dependency within *DNMT3A*-mutant LSC-like populations and heightened interferon-mediated responses ^64-66^. Clinical studies support this with non-inferior, and in some cohorts, even an improved rate of complete remission (CR; <5% marrow blasts) in *DNMT3A*-mutated *NPM1*^*c*^ AML patients treated with venetoclax-based regimens ^46,67-69^. Our data support these observations, and demonstrate that Ven+Aza combination effectively targets both NF and DNF LSCs and attenuates quiescence-associated transcriptional programs globally including in the deeply quiescent HSC-Like DNF LSCs. These results support early use of venetoclax-based combinations in *DNMT3A*^*mut*^*-NPM1*^*c*^*-FLT3*^*ITD*^ patients to achieve remission and facilitate progression to potentially curative allogeneic stem cell transplantation ^70,71^. More broadly, as the clinical use of venetoclax expands beyond patients unfit for intensive chemotherapy, our findings support a shift toward genetically-informed deployment of these regimens, with less reliance on age or fitness alone.

Although the differential chemotherapy resistance observed in DNF LSCs was substantially reduced by Ven+Aza treatment, a residual quiescent population persisted, with a relative enrichment of the rare MPP2 compartment in DNF AML. This suggests that therapy-induced remodeling of the LSC hierarchy as a potential mechanism of escape that warrants further investigation. These findings may explain why, despite improved initial remission rates, patients with triple *DNMT3A*^*mut*^*-NPM1*^*c*^*-FLT3*^*ITD*^ AML continue to exhibit inferior long-term outcomes and relapse following venetoclax-based therapy ^17,68^.

These findings highlight the need for therapeutic strategies that directly target LSC quiescence in *DNMT3A-*mutated AML. Interferon-α is known to promote HSC cell-cycle entry ^57,72^ and its long-acting formulation, pegIFNα, is currently used for treatment of myeloproliferative neoplasms (MPN) ^73^. Here, pegIFNα treatment in DNF AML prior to chemotherapy resulted in enhanced depletion of quiescent LSCs and marked contraction of the otherwise resistant LSC compartment. Of note, pegIFNα treatment alone was insufficient to significantly reduce G0 LSCs at 48 hours, likely reflecting the transient and dynamic nature of interferon-induced cell-cycle entry ^57,72^. These findings suggest that further optimization of timing and dosing of pegIFNα relative to chemotherapy maybe needed to exploit the full extent of LSC activation. Importantly, these data demonstrate that *Dnmt3a*^*R878H*^ induced quiescence is a reversible and therapeutically targetable state. By forcing cell-cycle entry, chemotherapy sensitivity in *Dnmt3a*-mutated AML can be restored.

Emerging data suggest that FLT3 inhibition may be particularly beneficial in *DNMT3A*^*mut*^*-NPM1*^*c*^*-FLT3*^*ITD*^ AML, with improved MRD clearance and 2-year overall survival reported ^74^. Notably, this murine FLT3^ITD^ model harbors the FLT3 F692L gatekeeper mutation, which confers resistance to all approved FLT3 inhibitors and has also been observed in patients following FLT3 inhibitor therapy ^75,76^. This limits our ability to access first- and second-generation FLT3i in this model, but also highlights the importance of evaluating third-generation FLT3 inhibitors, designed to overcome this resistance ^77^. Future work targeting NPM1-gene expression, for example, with menin inhibitors may represent a promising complementary strategy in *NPM1*^*c*^ AML ^78-80^. This may be particularly relevant in *DNMT3A*-mutant disease given the preferential activation of HOX-driven stemness programs ^27,81,82^. Rational combination strategies targeting both LSC quiescence and survival pathways may enable more effective eradication of *DNMT3A*-mutant LSCs and reduce risk of relapse. However, it is unknown whether *DNMT3A* mutations influence sensitivity to menin inhibition.

Collectively, our study identifies *Dnmt3a*^*R878H*^ mutation as a key epigenetic driver of LSC quiescence underlying chemotherapy resistance and disease persistence in *Npm1*^*c*^*-Flt3*^*ITD*^ AML. We demonstrate that a quiescent HSC-like LSC compartment represents a critical reservoir for disease persistence and relapse that is not fully eradicated by current therapies. Targeting LSC quiescence through combined cell-cycle activation with cytotoxic or targeted therapies therefore represents a promising strategy to achieve durable disease control.

## Acknowledgments

The *Dnmt3a*^*R878H*^ allele was generously provided by Prof. Marco J. Herold (CEO of the Olivia Newton-John Cancer Research Institute, Melbourne). Azacitidine was kindly provided by Bristol Myers Squibb (BMS) and Daniel Lopes de Menezes. Murine pegIFNα was gifted from PharmaEssentia. The authors gratefully acknowledge the technical support that was provided by staff at the QIMR Berghofer Animal Facility, Flow Cytometry Facility and Sequencing Facility. The valuable contributions and perspectives of the consumers and patients involved in this study are also gratefully acknowledged. J.S was supported by a Cancer Council Queensland fellowship (2025829). This work was supported by Cancer Australia’s Priority-driven Collaborative Cancer Research Scheme (PdCCRS, 2023/PCRS/0214; P.TS), the National Health and Medical Research Council (NHMRC, 1195987; S.W.L), the Gordon and Jessie Gilmour Leukaemia Research Trust and the Herron Family Trust.

## Contribution

P.TS conceptualized and designed the experiments, analyzed and interpreted the data, and wrote the manuscript. J.S conceptualized and design bioinformatics workflows, performed bioinformatic analyses and contributed to data interpretation and visualization. S.W.L and M.J.B, contributed to experimental conceptualization and design, data interpretation intellectual input, and study supervision. V.L contributed to data interpretation and clinical input. P.TS, E.C, S.H.N.C, S.A, R.H, Y.J and L.C performed the experiments and collected data. All authors contributed to and edited the manuscript.

## Data Availability

The raw and processed sequencing data generated in this study will be deposited in ArrayExpress under accession number XXX, including Dox+AraC RNA-seq (XXX), Ven+Aza RNA-seq (XXX), ATAC-seq (XXX), EM-seq (XXX), Dox+AraC 10x GeneFlex (XXX), and Ven+Aza 10x GeneFlex (XXX).

