## Supplementary Methods for "*Dnmt3a*-mutant Leukemia Stem Cells evade chemotherapy through enforced quiescence in *Npm1^c^-Flt3^ITD^* Acute Myeloid Leukemia"

**Mouse models and breeding.** All animal breeding and procedures were approved by the QIMR Berghofer Institutional Animal Ethics Committee (protocol A2109-620). Mice were maintained at the QIMR Berghofer animal facility under controlled conditions, including an 8-hour light/dark cycle, a temperature of 19-21 °C, and 55-65% humidity.

The conditional *Npm1*$^{flox\text{-}cA}$ knock-in and constitutively active *Flt3*$^{ITD}$ alleles were obtained from Prof. George Vassiliou [1] and Prof. Gary Gilliland [2], respectively. The constitutive *Dnmt3a*$^{R878H}$ allele, equivalent to human R882H, was generated by the Prof. Marco Herold, (MAGEC laboratory, The Walter and Eliza Hall Institute (WEHI)) as recently been described [3]. The *Rosa26*$^{creER}$ allele was obtained from The Jackson Laboratory (Strain #:008463).

Rosa26$^{CreER/CreER}$-*Npm1*$^{flox\text{-}cA/+}$ mice were crossed with mice homozygous for *Flt3*$^{ITD}$ to generate *Rosa26-Cre*$^{+/-}$-*Npm1*$^{cA/+}$-*Flt3*$^{ITD/+}$ animals (referred to as NF). To generate the triple-mutant *Rosa26-Cre*$^{+/-}$-*Dnmt3a*$^{R878H/+}$-*Npm1*$^{cA/+}$-*Flt3*$^{ITD/+}$ mice (referred to as DNF), *Rosa26-Cre*$^{+/+}$-*Npm1*$^{flox\text{-}cA/flox\text{-}cA}$ mice were first crossed with male *Dnmt3a*$^{R878H/+}$ mice, due to reported breeding difficulties in female *Dnmt3a*$^{R878H/+}$ mutants [3]. The resulting litters were subsequently crossed with *Flt3*$^{ITD/ITD}$ mice. All strains were maintained on a C57BL/6 background (CD45.2$^{+}$).

Genotyping for the *Rosa26-Cre*$^{+/-}$, *Npm1*$^{cA/+}$ and *Flt3*$^{ITD/+}$ was performed by PCR as previously descried [1, 2, 4]). *Dnmt3a*$^{R878H/+}$ mutant allele was genotyped utilising a SNP KASP assay (Cat# LGC-KBS-1050-121, GENEWORKS) as per manufacturer protocol, using allele-specific primers Allele X (5’-CAGTCTCTGCCTCGCCAAGC-3’), Allele Y (5’-GCAGTCTCTGCCTCGCCAAGT-3’) and common primer (5’-GTGTTTGGCTTCCCCGTCCACTA-3’) with analysis performed on ABI Quantstudio 5 qPCR system. The presence of *Dnmt3a*$^{R878H}$ was further confirmed via sanger sequencing using BigDye Terminator (v3.1; Thermo Fisher Scientific, Cat#4337455).

*Ptprc*$^{a/b}$ (CD45.1+/CD45.2+) congenic mice used for transplantation studies were generated from an F1 cross between wildtype C57BL/6J (Ptprc$^{b/b}$, CD45.2$^{+}$) and Ptprc$^{a/a}$ (CD45.1$^{+}$) mice on a C57BL/6J background. Wildtype C57BL/6J and Ptprc$^{a/a}$ C57BL/6J mice were purchased from the ARC Animal Resources Centre or Ozgene (WA, Australia).

**Peripheral Blood Analysis.** Peripheral blood was collected via the submandibular vein into EDTA-coated tubes (Greiner Bio-One, Cat# 450532). Complete blood counts were analysed using either a Hemavet 950 analyser (Drew Scientific) or an Element HT5 analyser (Heska). Peripheral blood smears were prepared and stained with Wright–Giemsa stain (BioScientific) according to manufacturer’s protocol.

For monitoring AML engraftment, 30 µL of peripheral blood samples were subjected to red blood cell lysis using BD Pharm Lyse™ Lysing Buffer (BD Biosciences, Cat# 555899). Cells were then stained with antibodies against mouse CD45.1-AF700 (A20, Biolegend, Cat# 110724) and CD45.2-V500 (donor AML; B10.S, BD Horizen™, Cat#562129) to assess donor-derived leukemic engraftment, along with additional markers as specified for each experimental panel. Samples were analysed by flow cytometry using a BD LSRFortessa (BD Biosciences).

**Bone marrow and Spleen harvest.** Bone Marrow (BM) cells were isolated by flushing the marrow from the hind limbs (femurs, tibias, and pelvis) through a 70 µm cell strainer (Greiner Bio-One, Cat# 4552) using a syringe fitted with a 26-gauge needle containing cold PBS supplemented with 2% foetal bovine serum (FBS; Thermo Fisher Scientific, Cat# 10099141). Splenocytes were isolated by mechanically dissociating the entire spleen through a 70 µm cell strainer using cold PBS supplemented with 2% FBS. Red blood cells in each tissue were then lysed (BD Biosciences, Cat# 555899). Viable BM and spleen cell suspensions were enumerated using trypan blue exclusion (Thermo Fisher Scientific, Cat#15250061) with an automated cell counter (Corning).

**Colony Forming Unit assay.** BM cells harvested from DNF and NF mice with overt AML were resuspended at 30,000 cells in 3 mL of MethoCult M3234 (STEMCELL Technologies, Cat# 03234) supplemented with murine stem cell factor (mSCF; 50 ng/mL, Miltenyi Biotec, Cat# 130-101-697), murine interleukin-3 (mIL-3; 10 ng/mL, PeproTech, Cat# 213-13), murine interleukin-6 (mIL-6; 10 ng/mL, PeproTech, Cat# 216-16), erythropoietin (EPO, 300U; PeproTech, Cat#100-64), and Penicillin–Streptomycin (1:100, Thermo Fisher Scientific, Cat# 15140122) and plated in triplicates. Colonies were counted after 7 days of culture, and cells were subsequently passaged weekly in fresh methylcellulose medium for a total of five passages to assess serial replating capacity.

**Drug Treatments.** For short-cycle standard chemotherapy studies, DNF and NF mice were stratified on day 18 post-transplantation to receive cytarabine (AraC; Pfizer) and doxorubicin (Dox; Pfizer) or vehicle saline control. Stock solutions of AraC (1 g in 10 ml isotonic water) and Dox (50 mg in 25 ml saline) were freshly diluted in saline (0.9% sodium chloride for irrigation; Thermo Fisher Scientific) prior to administration. Final dosing concentrations were 50 mg/kg body weight for AraC and 1.5 mg/kg body weight for Dox in a total injection volume of 200 µl per recipient mouse. AraC and Dox were co-administered intravenously via tail vein injection on days 1–3, as previously described [5, 6].

To model a shortened treatment cycle of venetoclax and azacitidine based on the clinical 28-day regimen, DNF and NF mice were stratified on day 12 post-transplantation to receive single-agent venetoclax (Ven; MedChem Express, Cat#HY-15531-5G), azacitidine (Aza; Bristol Myers Squibb), or the Ven+Aza combination. Drugs were administered on days 1–5, followed by days 8 and 9 (5+2 regimen). Ven was administered via oral gavage with a dose ramp-up schedule of 25% on day 1, 50% on day 2, and 100% from day 3 onwards to a final dose of 100 mg/kg body weight. Venetoclax was prepared in vehicle consisting of 5% dimethyl sulfoxide (DMSO; Sigma Aldrich), 40% PEG300 (MedChem Express, Cat#HY-Y0873-100ML), 5% Tween-80 (MedChem Express, Cat#HY-Y1891-100ML), and 50% saline. Aza was freshly prepared in saline and administered Intraperitoneally within 2 hours of preparation at 2 mg/kg body weight. Control mice received 200 µl of vehicle solution consisting of 5% DMSO, 40% PEG300, 5% Tween-80, and 50% saline.

To induce cell cycle entry in $Dnmt3a^{R878H}$-mutant LSCs, DNF mice received a single dose of 600 ng ropeginterferon-α11 (pegIFNα; mP1101, PharmaEssentia) administered subcutaneously in 200 µl of 0.1% bovine serum albumin (BSA) in phosphate-buffered saline (PBS). Treatment was performed on day 17 post-transplantation following stratification, and mice were analysed 48 h after pegIFNα administration.

For pegIFNα and chemotherapy combination studies, mice were first primed with pegIFNα treatment. Forty-eight hours later, mice received three doses of Dox/AraC chemotherapy as described above. Mice were analysed 24 h after completion of the final chemotherapy dose.

**Flow cytometry analysis.** Surface staining for flow cytometry was performed on erythrocyte-lysed single-cell suspensions derived from BM, spleen, or peripheral blood. Cells were stained in PBS supplemented with 2% FBS for 30 min at 4 °C. All antibodies are used at 1:100 dilution and purchased from Biolegend unless otherwise specified.

AML donor cells were distinguished from recipient cells using anti-mouse CD45.1-AF700 and CD45.2-V500 or CD45.2-PerCP-Cy5.5 (B10.S, BD Horizon™, Cat# 562129) antibodies, as described above. For mature AML cell phenotyping, cells were stained with antibodies against CD45.1-AF700, CD45.2-V500, CD3-PE (17A2, Cat# 100206), B220-PerCpCy5.5 (RA3-6B2, Cat# 103236), Mac1-APC-Cy7 (M1/70, Cat# 101226), Gr1-AF488 (RB6-8C5, Cat# 108417), and c-Kit-APC (2B8, Cat# 105812).

For phenotyping of the haematopoietic stem and progenitor cell (HSPC) compartment, $3 \times 10^6$ BM cells were first labelled with a biotin-conjugated lineage antibody cocktail consisting of Ter119 (TER-119, Cat# 116204), Gr1 (RB6-8C5, Cat# 108404), Mac1 (M1/70, Cat# 101204), B220 (RA3-6B2, Cat# 103204), CD3e (145-2C11, Cat# 100304), and CD5 (53-7.3, Cat# 100604). Cells were subsequently stained with BV605-conjugated streptavidin (Cat# 405229). HSPC populations were further resolved using c-Kit-APC, Sca1-PE-Cy7 (D7, Cat# 108114), CD16/32 (FcγR; 93, Cat# 101328), CD34-FITC (RAM34, Thermo Fisher Scientific, Cat# 11-0341-82; at 1:50 dilution), CD48-PE (HM48-1, Cat#

103406), and CD150-PE-Cy5 (TC15-12F12.2, Cat# 115912). Cells were washed with PBS + 2% FBS and resuspended in SYTOX Blue Dead Cell Stain (Thermo Fisher Scientific, Cat# S34857) for dead-cell exclusion.

For cell cycle analysis, biotin-labelled BM cells were depleted using BD IMag Streptavidin Particles Plus (BD Biosciences, Cat# 557812). Remaining cells were normalised to 3 × $10^6$ cells per sample and stained with surface markers to identify AML MPP3 cells (CD45.1-AF700, CD45.2-PerCP-Cy5.5, streptavidin-APC-Cy7 (Calt#405208), c-Kit-APC, Sca1-PE-Cy7, CD48-PE, and CD150-PE-Cy5). Following surface staining, cells were fixed and permeabilised using the Fix and Perm Cell Permeabilization Kit (Invitrogen, Cat# GAS004) and stained with Ki-67-AF488 (BD Pharmingen™, B56, Cat# 561165) according to the manufacturer's instructions. Cells were subsequently washed and resuspended in PBS + 2% FBS containing 0.2 mg/ml Hoechst 33342 (Invitrogen, Cat# H3570) for DNA content analysis.

For Fluorescence Activated Cell Sorting (FACS) of LSK and MPP3 populations, lineage-depleted BM cells were stained with the HSPC antibody panel described above. Flow cytometric analysis and sorting were performed using a BD LSRFortessa or BD FACSAria (BD Biosciences) with FACSDiva software. Post-acquisition analysis was conducted using FlowJo v10.7.2 (Treestar, CA). Data from gated populations were calculated from a minimum of 100 events.

**RNA sequencing sample processing and analysis.** For baseline and chemotherapy-treatment experiments, 20,000 MPP3 cells per genotype and treatment condition were isolated using a BD FACSAria cell sorter and sorted directly into 100 µL of Extraction Buffer from the Arcturus PicoPure RNA Isolation Kit (Thermo Fisher Scientific, Cat# KIT0204). Total RNA was extracted according to the manufacturer's protocol, including on-column DNase treatment to remove genomic DNA contamination. Oligo d(T)-captured mRNA (10 ng input RNA per sample) was processed for next-generation sequencing (NGS) library preparation using the NEBNext® Ultra™ II RNA Library Prep Kit for Illumina (New England Biolabs, Cat# E7770S) according to the manufacturer's protocol. Poly(A)+ mRNA was enriched using the NEBNext Poly(A) mRNA Magnetic Isolation Module (New England Biolabs, Cat# E7490S). cDNA purification was performed using SPRIselect Reagent Kit (Beckman Coulter, Cat# B23317). Libraries were dual indexed using NEBNext Multiplex Oligos for Illumina (96 Unique Dual Index Primer Pairs) (New England Biolabs, Cat# E6440S) with 15 cycles of PCR amplification, as recommended by the manufacturer. RNA sequencing was conducted across 4 independent biological leukaemia replicates per genotype and treatment.

For Venetoclax and Azacitidine (Ven+Aza) studies, total RNA was extracted from up to 20,000 sorted MPP3 cells per sample using the Arcturus PicoPure RNA Isolation Kit (Thermo Fisher Scientific, Cat# KIT0204) as described above. One nanogram of total RNA per sample was used for cDNA synthesis and library preparation using the SMART-Seq® mRNA LP Kit (Takara Bio, Cat# 634768) according to the manufacturer's instructions. Briefly, poly(A)+ mRNA was reverse transcribed using an oligo-d(T) primer, and full-length cDNA was generated using SMART (Switching Mechanism at 5′ end of RNA Template) template-switching technology followed by PCR amplification. Libraries were indexed using the SMART-Seq mRNA LP Unique Dual Index Kit (1–24) (Takara Bio, Cat# 634756). RNA sequencing was conducted on two independent biological leukaemia samples, each consisting of two technical replicates per genotype and treatment condition.

For all the experiments, RNA concentration was first quantified using the Qubit RNA High Sensitivity (HS) Assay Kit (Invitrogen, Cat# Q32852). RNA quality and integrity were subsequently assessed using the Agilent High Sensitivity RNA ScreenTape assay (Cat# 5067-5580) for TapeStation system. All samples exhibited RNA integrity numbers (RIN) between 9.5 and 10. Amplified Library fragment size, quality and quantity was validated using the Agilent 2100 Bioanalyzer and Agilent's High Sensitivity D5000 Kit (Cat# 5067-5593). Final libraries were pooled and sequenced on an Illumina NextSeq 2000 platform using a High Output P2 flow cell with paired-end 200 bp reads, generating 400 million reads per run (25M reads/sample). Sequencing was performed using NextSeq System Suite (version 2.1.2).

Reads were trimmed for adapter sequences using Cutadapt v1.11 and aligned using STAR v2.5.2a to the GRCm38 genome build, with assembly using the gene, transcript, and exon features of Ensembl (release 67). Expression was estimated using RSEM v1.2.30, with transcripts with zero read counts across all samples removed prior to analysis. Normalisation and the differential expression analysis was performed using edgeR. Gene set enrichment analysis (GSEA) was performed using Broads GSEA v 4.1.0 or fgsea R package on logFC of genotype of treatment differences of interest. Gene sets assessed were obtained from MSigDB collections, including Hallmark and C2, or are listed in Supplemental Table 1.

**ATAC-Sequencing and analysis.** The Assay for Transposase-Accessible Chromatin using sequencing (ATAC-seq) was performed using the ATAC-Seq Kit (Active Motif, Cat# 53150) according to the manufacturer's protocol (version B9). Briefly, 50,000 MPP3 cells per sample from independent biological DNF and NF leukaemia samples, each consisting of four technical replicates per genotype, were FACS-sorted into PBS + 2% FBS, washed, and lysed in ATAC lysis buffer to isolate nuclei. The nuclei were subsequently subjected to Tn5 transposase-mediated tagmentation. Tagmented DNA was PCR-amplified using unique i7 and i5 index primers provided with the kit. The resulting libraries were quantified and assessed for quality using the D1000 DNA ScreenTape assay (Agilent Technologies, Cat# 5067-5583). Final libraries were pooled and sequenced on an Illumina NextSeq 2000 platform using a High Output P2 flow cell with paired-end 100 bp reads (400M; 25M reads/sample). Sequencing was performed using NextSeq System Suite (version 2.1.2).

Raw reads were mapped with bwa mem to the GRCm38 genome build. Duplicated reads were marked with picard v 2.18.15. Supplementary alignments and poor quality read pairs were filtered with -F 3852. Read depth by sequence coordinates in wig format were generated with samtools v1.17 mpileup and transformed to bigwig with wigToBigWig as implemented in homers perl v5.28.0 module. For each mapped bam file peaks were called in python v2.71.10 with macs2. Bedops v2.4.35 was used to generate the union of all individual sample's peak coordinates. Bedtools v2.27.1 coverageBed option was used to extract the read coverage under the peak for each sample. The matrix of peak read coverage by sample was used for downstream analysis. Peaks were annotated using annotatePeak function from the ChIPseeker R package v1.44.0 with TSS region transcription -2000 and 1000 with TxDB TxDb.Mmusculus.UCSC.mm10.knownGene v3.10.0 and annoDb "org.Mm.eg.db". Differential peak analysis was performed with edgeR R package and p-value adjustment using false discovery rate.

**Methylation sequencing and analysis.** Genome-wide DNA methylation profiling of LSCs was performed using 50,000 FACS-sorted MPP3 cells obtained from independent DNF and NF leukaemia samples (plus two technical replicates per genotype). Genomic DNA was isolated using the QIAamp DNA Micro Kit (Qiagen, Cat# 56304). Approximately 20 ng of genomic DNA per sample was supplemented with unmethylated lambda DNA and CpG-methylated pUC19 DNA spike-in controls and fragmented using a Covaris S220 sonicator (Covaris) with the following parameters: peak power 150, duty factor 10%, 200 cycles per burst for 70 seconds, yielding DNA fragments of approximately 350–400 bp. Fragment size distribution was confirmed using the High Sensitivity D1000 DNA ScreenTape assay (Agilent Technologies, Cat# 5067-5583). Whole methylome analysis was performed using the NEBNext® Enzymatic Methyl-seq Kit (New England Biolabs, Cat# E7120S) according to the manufacturer's instructions. Briefly, fragmented DNA underwent end repair and dA-tailing followed by ligation of EM-seq adapters. Libraries were then subjected to enzymatic cytosine conversion reactions that selectively convert unmethylated cytosines, while preserving 5-methylcytosine (5mC) and 5-hydroxymethylcytosine (5hmC). DNA denaturation was performed using formamide (Sigma-Aldrich, Cat# F9037). Converted DNA libraries were PCR-amplified using NEBNext Low Volume Unique Dual Index primers, pooled, and sequenced by Novogene on a NovaSeq X Plus platform using paired-end 150 bp reads, targeting approximately 98 Gb of raw sequencing data per sample.

Reads were mapped using bwa-meth (https://github.com/brentp/bwa-meth) with default parameters. MethylDackle (https://github.com/dpryan79/MethylDackel) was used to calculate the percentage of methylated C in a CpG context. methylDackle bedgraphs were sorted with bedtools and converted to bigwig using bedgraphToBigWig function as implemented in homer v4.8. Differential methylation analysis was performed with the methylKit R package v1.34.0. CpGs were annotated using annotatePeak function from the ChIPseeker R package v1.44.0 with TSS region transcription -2000 and 1000 with TxDB TxDb.Mmusculus.UCSC.mm10.knownGene v3.10.0 and annoDb "org.Mm.eg.db".

**Single Cell gene-expression profiling.** DNF and NF LSK populations were FACS-sorted following treatment with Dox/AraC or Ven+Aza, alongside their corresponding vehicle-treated controls. For standard chemotherapy studies, ScRNA-seq was performed on three independent DNF and one NF leukaemia samples. For venetoclax-based studies, sequencing was performed on four independent DNF and three NF leukaemia samples. Cells were fixed using the Chromium Next GEM Single Cell Fixed RNA Sample Preparation Kit (10x Genomics, Cat#PN-1000414) according to the manufacturer's instructions (CG000478, Rev D). Due to variable depletion of the LSK population following treatment in both DNF and NF samples, lineage-depleted BM cells from recipient mice of each individual leukaemia were pooled (up to 12 mice per genotype per treatment) prior to FACS sorting to obtain the recommended input for the fixed RNA workflow. Sorted cell numbers ranged from 100,000 to 400,000 cells per sample. Cells were fixed in fixation buffer containing formaldehyde (37%; Thermo Fisher Scientific, Cat# BP531-25) for 20 hours at 4°C, followed by quenching. Fixed cells were supplemented with a final concentration of 10% glycerol (Merck Life Science, Cat# G5516-100ML) and stored at -80°C between separate experiments until further processing.

Single-cell RNA expression profiling of fixed cells was performed using the Chromium Fixed RNA Kit Mouse Transcriptome (4 × reactions × barcodes; 10x Genomics, Cat# PN-1000496) according to the manufacturer's instructions. The Chromium Fixed RNA workflow allows processing of four individual samples per reaction using probe-based multiplexing. In the initial sequencing run, samples were multiplexed using the four probe sets provided in the kit, allowing conditions such as DNF Dox/AraC, NF Dox/AraC, DNF vehicle, and NF vehicle to be barcoded individually and processed within the same reaction. In subsequent experiments, to increase the number of biological replicates analysed, additional multiplexing using sex-mismatched samples was implemented. Upon recovery after storage, equal cell numbers from one male and one female leukaemia sample of the same genotype (vehicle- or drug-treated) were combined and processed as a single sample for downstream probe hybridization. Briefly, cells were incubated with mouse Whole Transcriptome Analysis (WTA) probes for approximately 20 hours at 42°C for probe hybridization. Following hybridization, equal numbers of cells per probe reaction were pooled and processed according to the pooled wash workflow. Cell concentrations were determined using trypan blue exclusion and a haemocytometer, and sufficient cells were loaded to target 40,000 recovered cells per reaction for Gel Bead-in-Emulsion (GEM) generation using the Chromium Next GEM Chip Q Single Cell Kit (10x Genomics, Cat# PN-1000418).
Following GEM generation and reverse transcription, cDNA was recovered and purified using SPRIselect beads (Beckman Coulter, Cat# B23317). Sequencing libraries were indexed using the Dual Index Kit TS Set A (10x Genomics, Cat# PN-1000251). Final libraries were quantified and quality assessed using the High Sensitivity D1000 DNA ScreenTape assay (Agilent Technologies, Cat# 5067-5366). Libraries were pooled and sequenced on an Illumina NextSeq 2000 platform using a High Output P4 flow cell with paired-end 100-cycle sequencing, targeting 10,000 reads per cell. Sequencing was performed using NextSeq System Suite (version 2.1.2).

Chromium Mouse Transcriptome Probe Set v1.0.1. Reads were mapped against the mm10 genome and annotated against the transcriptome mm10-2020-A with cellranger v8.0.1. Data were analysed in R v4.5.0. Normalisation, filtering (mitochondrial % <5 and # RNA features >200) and cell cycle scoring was performed with Seurat v5.4.0. Integration was performed identifying integration anchors of the first 30 reciprocal PCA reduction. The 20 k-nearest neighbours on the first 20 integrated dimensions were identified and using to perform clustering with shared nearest neighbor (SNN) modularity optimization-based clustering algorithm and res=0.8. Clusters were annotated by consensus call from known marker gene expression, performing ssGSEA (GSVA R package v2.2.1) against marker gene sets from sorted or single cell HSPC population ([7-9]; Supplemental Table 1) and Seurat mapping against the mouse bone marrow atlas [10] (downloaded from synapse.org with Synapse ID: syn66721894).

**Public data set analysis.** RNA-Seq data from de novo AML patients from Corces et al 2016 were downloaded from GEO (GSE74246)[11]. Lin– CD34+ CD38– TIM3– CD99– sorted HSCs from NF (SU048, SU209) and DNMT3A mutant NF (SU353, SU575) were extracted. The Leucegene AML cohort's RNA-Seq data were downloaded from GEO with GSE49642. Whole bone marrow data from patients annotated NF (03H116, 10H115, X07H158, X13H110) and DNMT3AR882H/C NF (X10H101, X10H092, X04H112, X09H002, X12H010) as indicated in the papers Supplemental Table1 were extracted [12]. Both datasets raw counts were normalised with edgeR. Normalised read counts were analysed for gene set enrichment comparing DNF vs NF patients with GSEA v4.1.1 custom pathways as listed in Supplemental Table 1. Pre-processed and annotated CD34+ single cell data from AML patients as generated by Beneyeto-Calabuig *et al 2023* [13] were downloaded from Figshare (https://doi.org/10.6084/m9.figshare.20291628; accessed 25/06/2024). Data from DNF patient A.8 and NF patient A.10 were extracted. Single cell GSEA was performed on normalised data with the GSVA R package v2.2.1 and default parameters. To assess differential gene expression between genotypes leukemic and immature annotated cells were extracted.
