## Supplemental Figures for "*Dnmt3a*-mutant Leukemia Stem Cells evade chemotherapy through enforced quiescence in *Npm1^c^-Flt3^ITD^* Acute Myeloid Leukemia"

### Supplemental Figure 1

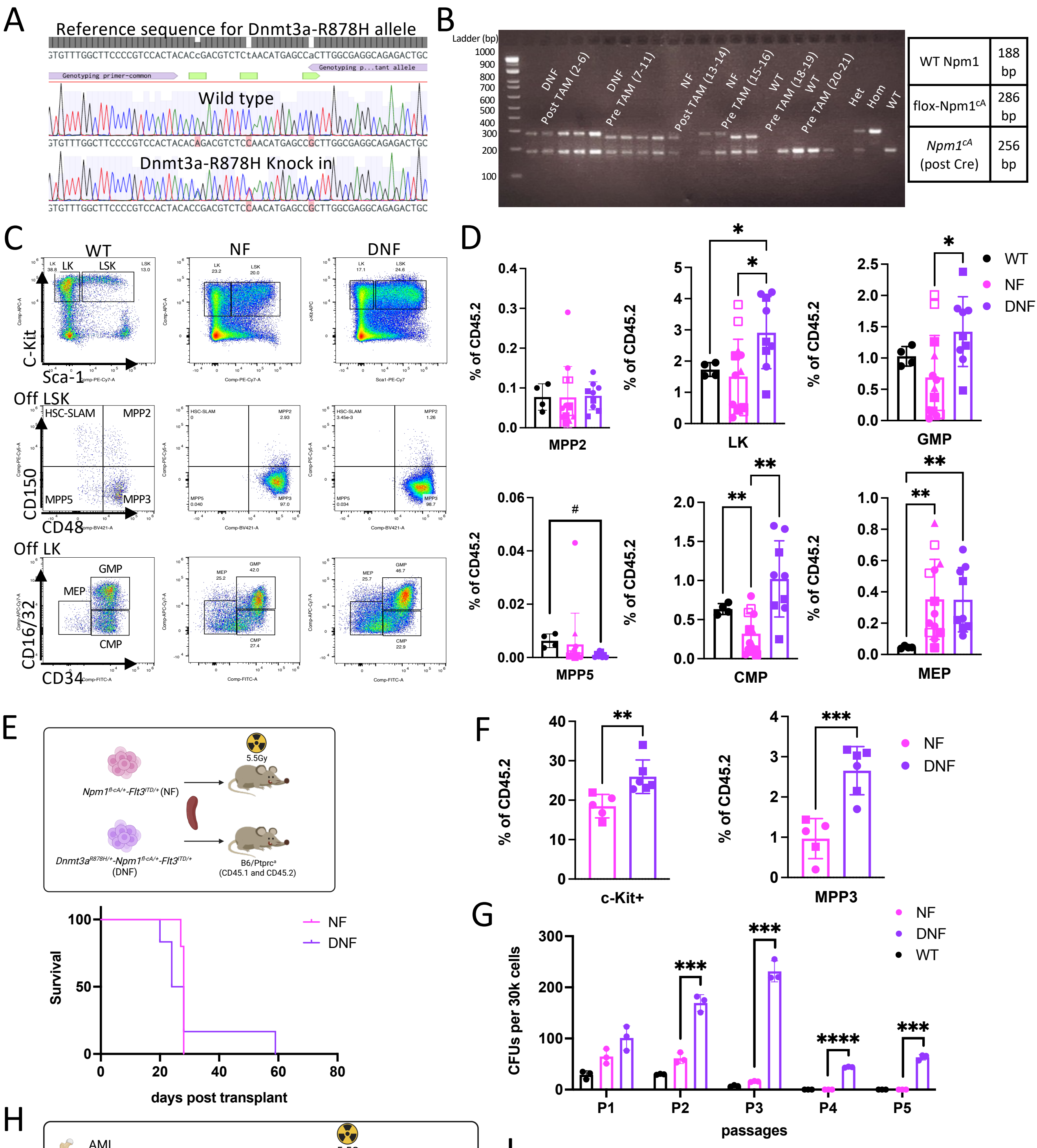


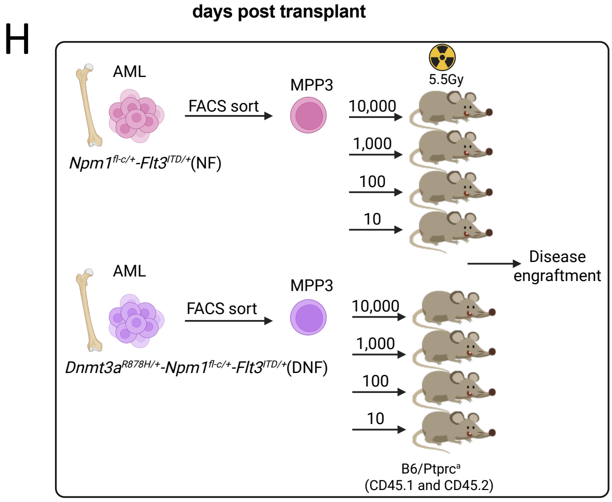


I

| Group | Dose | Number of animal tested | Number of animals that developed leukaemia |
|---|---|---|---|
| DNF | 10,000 | 9 | 8 |
| | 1000 | 9 | 6 |
| | 100 | 8 | 3 |
| | 10 | 9 | 0 |
| NF | 10,000 | 9 | 8 |
| | 1000 | 9 | 3 |
| | 100 | 9 | 3 |
| | 10 | 9 | 0 |

1/ (LSC frequency)

| Group | Lower | Estimate | Upper |
|---|---|---|---|
| DNF | 4207 | **1796** | 767 |
| NF | 9926 | **2703** | 1234 |

| Chisq | DF | P-value |
|---|---|---|
| 0.774 | 1 | 0.379 |

### Supplemental Figure 2

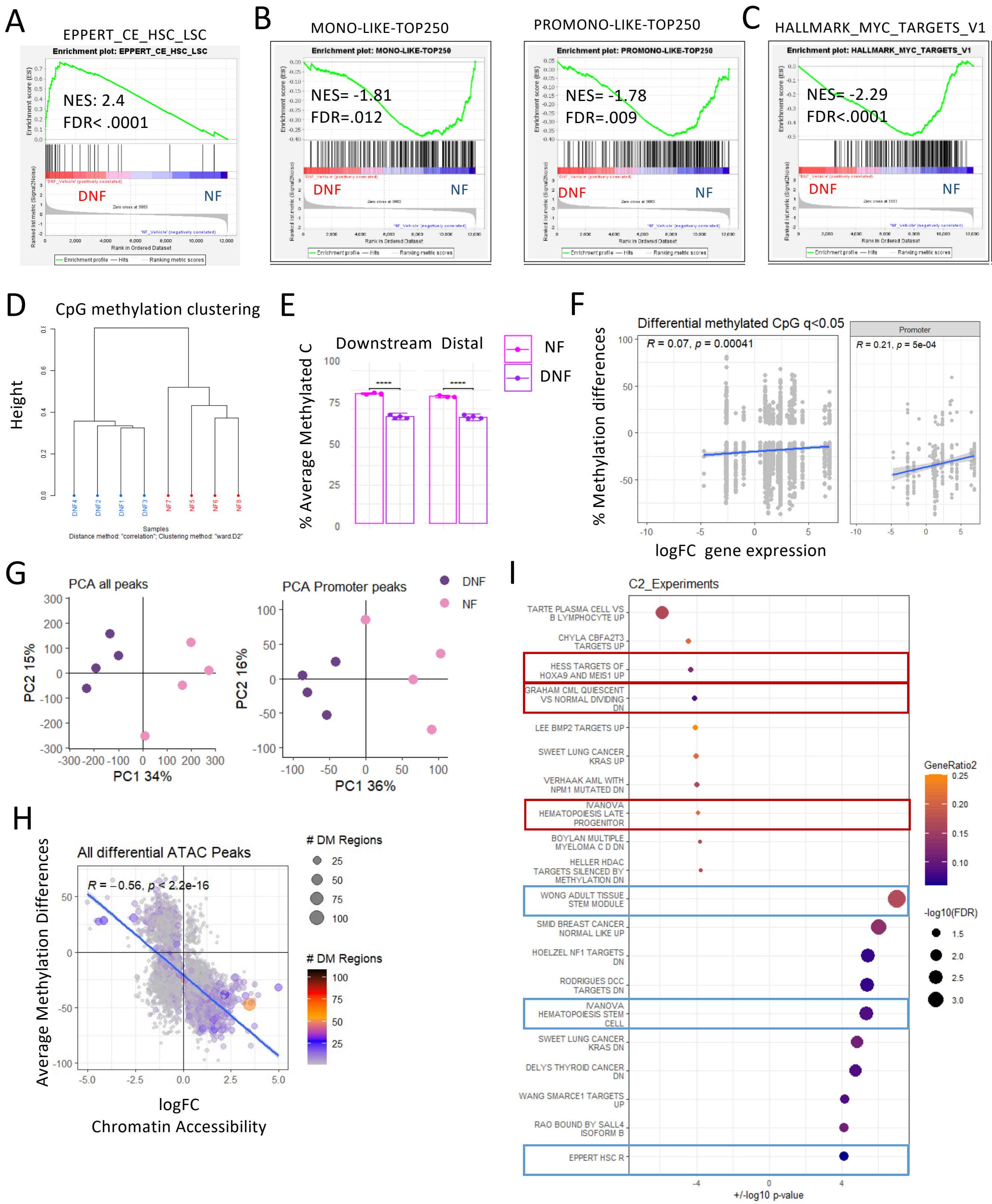

### Supplemental Figure 3

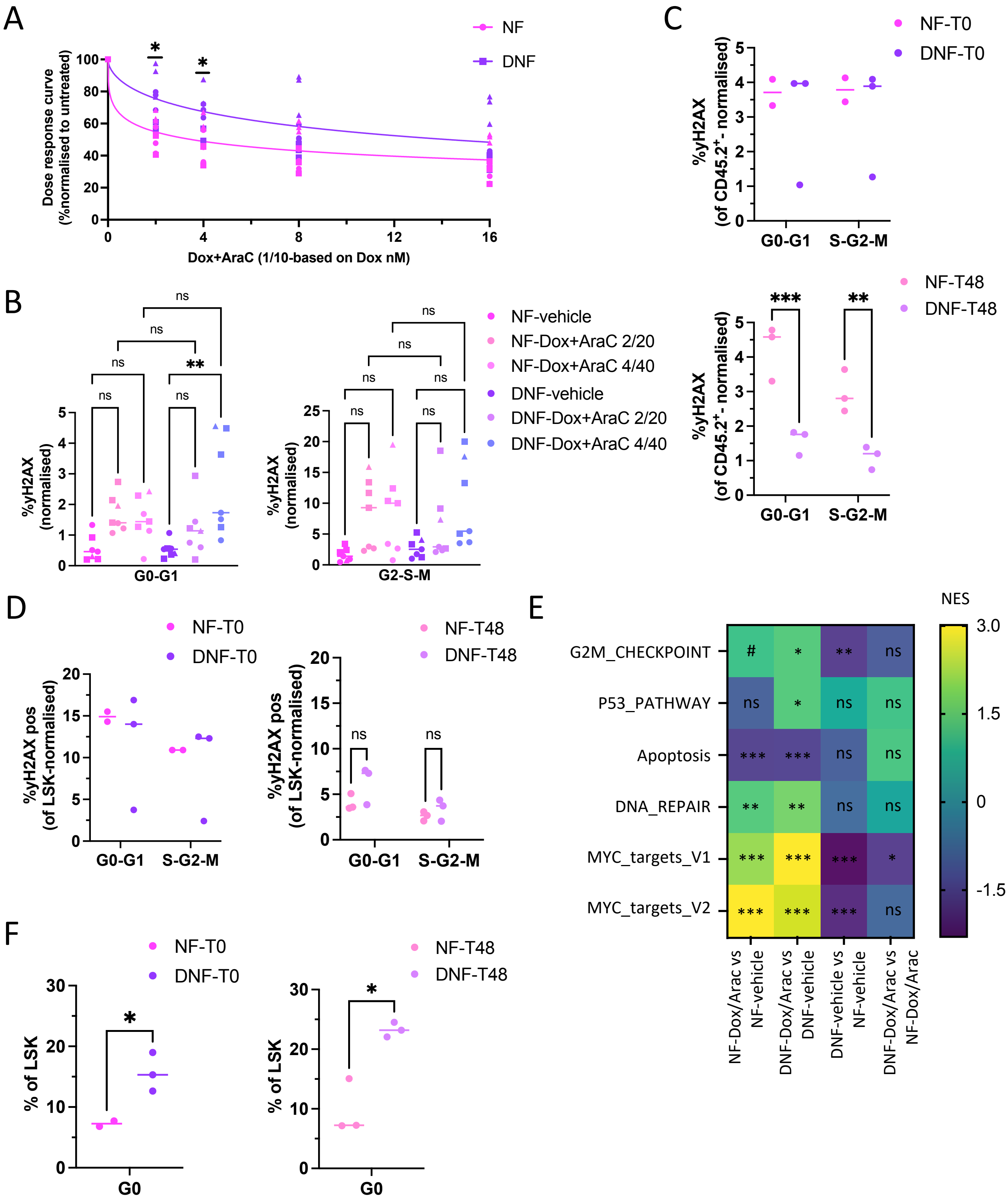

### Supplemental Figure 4

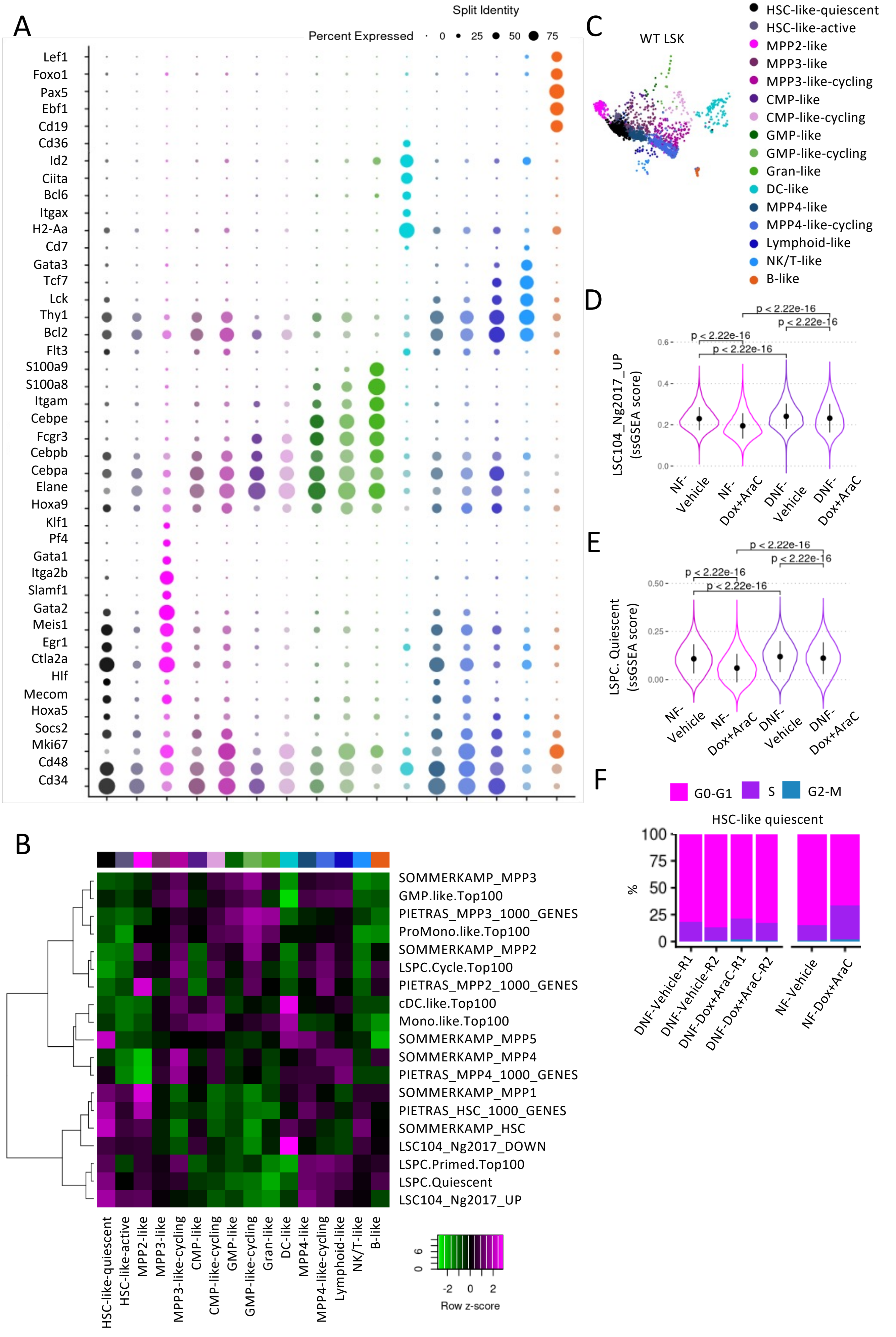

### Supplemental Figure 5

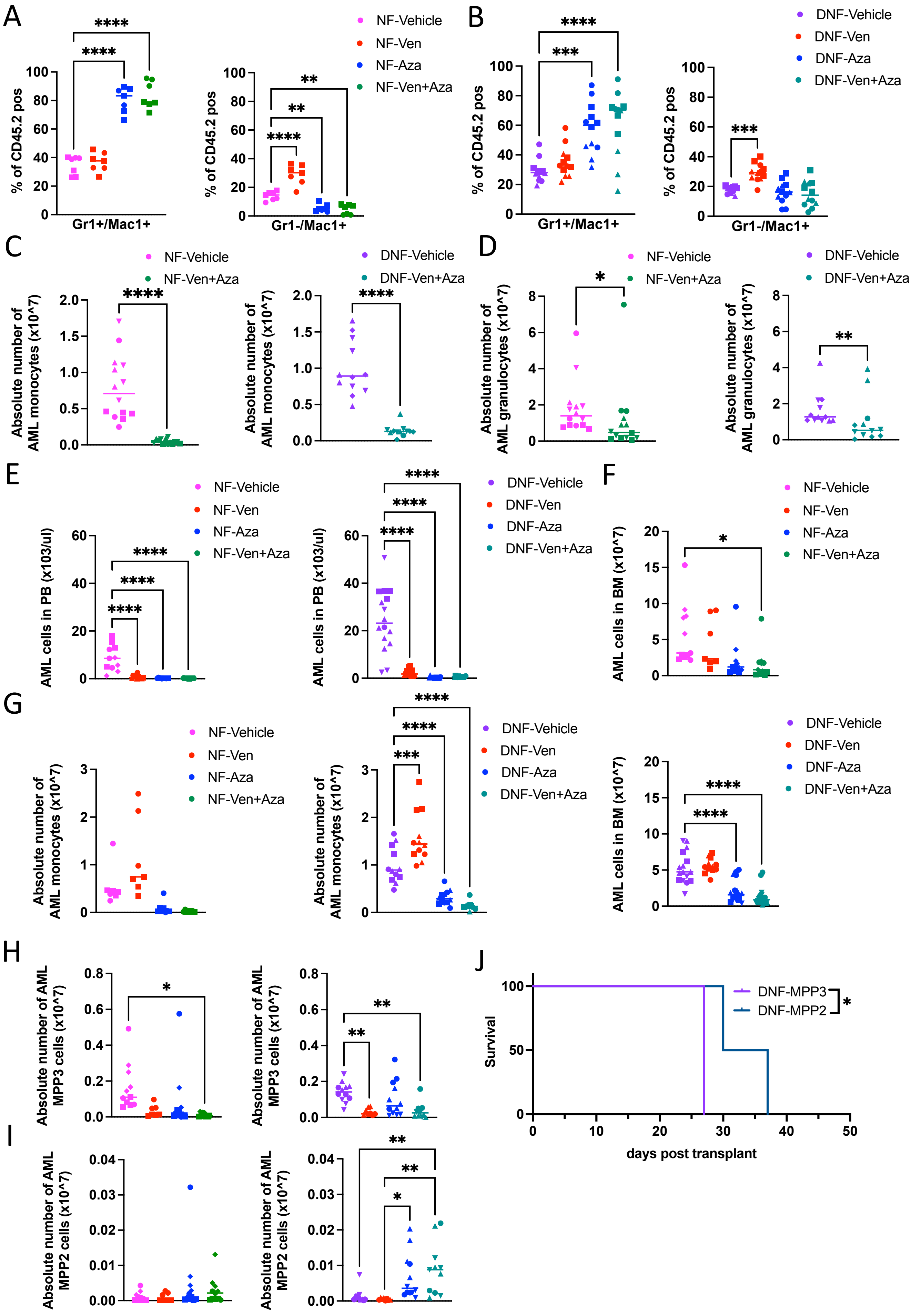

### Supplemental Figure 6

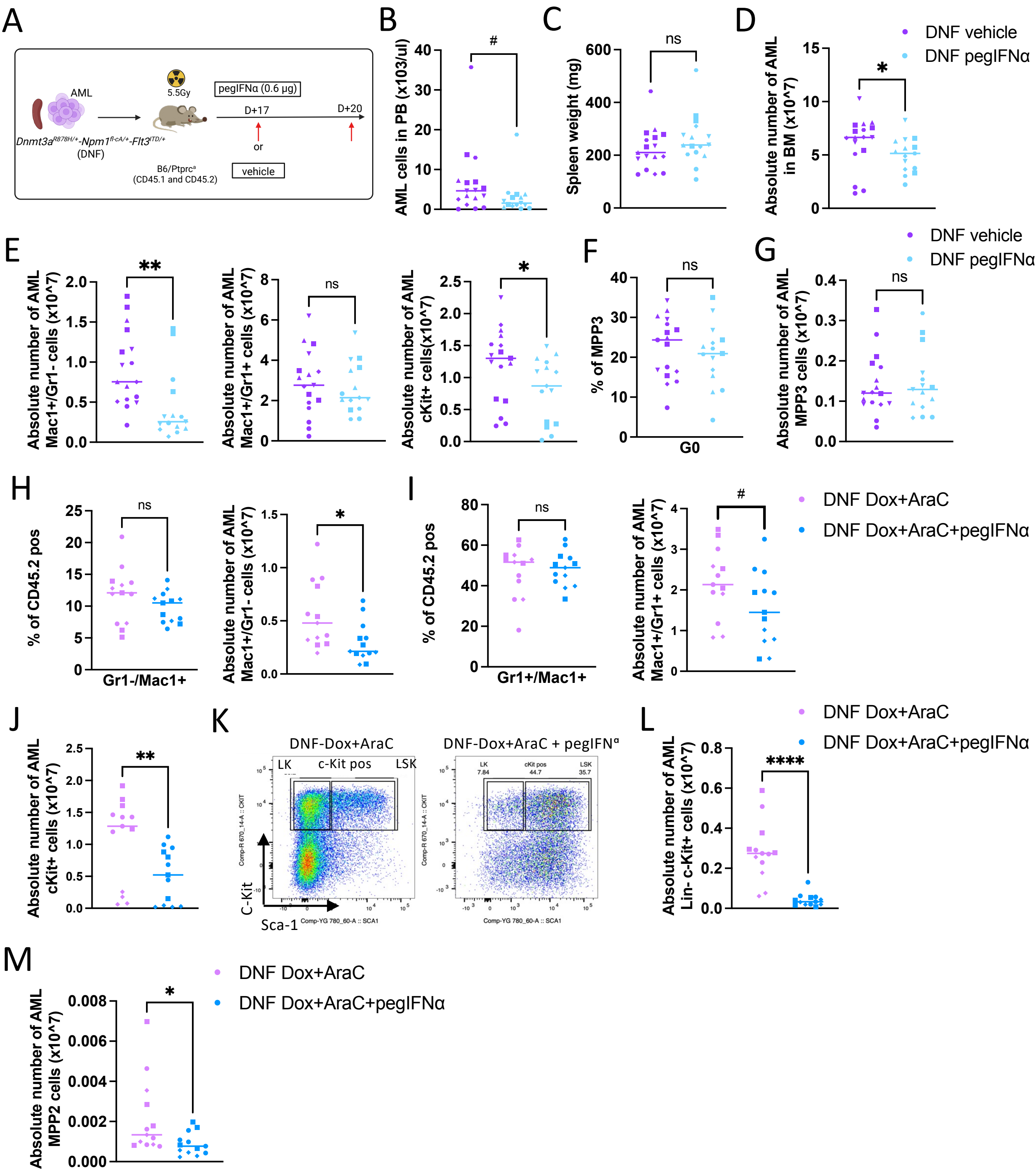
