## Supplemental Figure Legends for "*Dnmt3a*-mutant Leukemia Stem Cells evade chemotherapy through enforced quiescence in *Npm1^c^-Flt3^ITD^* Acute Myeloid Leukemia"

**Supplementary Figure 1. *Dnmt3a*$^{R878H}$ co-expression enhances expansion of the LSC-enriched MPP3 population but does not significantly increase functional LSC frequency within an equivalent number of MPP3 cells.** (A) Sanger sequencing chromatogram confirming *Dnmt3a*$^{R878H}$ knock in allele in DNF AML sample. (B) Pre- and post-TAM genotyping PCR performed on PB samples from DNF (lanes 2–6, post-TAM; 7–11, pre-TAM), NF (lanes 13–14, post-TAM; 15–16, pre-TAM), and WT controls (lanes 18–19, post-TAM; 20–21, pre-TAM). PCR primers were designed around *Npm1* to distinguish WT, floxed, and post-Cre (cA) alleles as previously described (1). (C) Representative flow cytometry plots showing the downstream gating strategy for LSK and LK populations in NF, DNF, and WT BM cells. (D) Flow cytometric analysis of populations within the HSPC compartment of primary AML BM (n= 4 WT, 13 NF, 9 DNF) (E) Experimental design and Kaplan-Meier survival analysis assessing the transplantability of NF and DNF AML into secondary recipient mice. (F) Flow cytometric quantification of c-Kit$^{+}$ and MPP3 population frequency in secondary AML BM (n= 5 NF, 6 DNF) (G) Colony-forming unit (CFU) counts following serial replating (5 weeks) of AML-derived cells from NF and DNF mice compared with WT controls. (H) Serial dilution transplantation assay in which mice received the indicated numbers of FACS-sorted MPP3s from NF or DNF AML. (I) Summary table showing the number of mice developing AML in each dilution/genotype cohort and the calculated LSC frequency within the LSC-enriched MPP3 population of NF and DNF mice. Each data point represents a biological replicate; identical symbols denote technical replicates derived from the same biological sample pooled from multiple experiments. Statistical analyses were performed using ordinary one-way ANOVA (D, G) or unpaired Student T-test with Welch' correction (F). Error bars represent mean ± SD. #p=0.06, *p<0.05, **p<0.01 and ***p<0.001.

**Supplementary Figure 2. Hypomethylated promoter regions in *Dnmt3a*$^{R878H}$-mutated LSCs are enriched for genes associated with HSC identity and stemness.** Bulk RNA-seq derived GSEA with NES and FDR of DNF (n=4) versus NF (n=4) LSCs for (A) HSC-associated gene signatures, (B) monocyte-like differentiation programs, and (C) MYC target gene signatures. (D) Hierarchical clustering of CpG methylation profiles in DNF and NF LSCs using correlation distance and Ward.D2 linkage. (E) Average percentage of DNA methylation across downstream, and distal regulatory regions in DNF and NF LSCs. (F) Gene expression changes (logFC) versus differences in DNA methylation at differentially methylated CpGs (FDR<0.05) in DNF compared to NF LSCs, including all differentially methylated CpGs (left) and promoter-associated CpGs (right). (G) Principal component analysis (PCA) of chromatin accessibility in DNF and NF LSCs using all ATAC-seq derived peaks (left) or promoter-associated peaks (right). Percentage of variance explained by each principal component (PC) is indicated on the axes. (H) Scatterplot comparing DNF versus NF differential ATAC peaks (FDR<0.05) as logFC change in chromatin accessibility versus differences in average DNA methylation under the ATAC peak (FDR<0.05). Point size and colour indicate the number of differentially methylated CpGs under each ATAC peak. R- and P-value are shown. (I) Dot plot showing hypergeometric enrichment of gene signatures associated with differentially methylated promoters in DNF versus NF LSCs (FDR<0.05). Log10 P-values are shown on the x-axis, with negative and positive values indicating signatures associated with methylation gain or loss in DNF, respectively. Only significantly enriched signatures are shown (p<0.05). Red and blue boxes highlight stemness/quiescence-associated and differentiation-associated gene signatures, respectively. Dot size represents −log10(FDR), and colour indicates gene ratio. (F, H) Display coefficients (R) and p-values of a Pearson's correlation test. Error bars represent mean ± SD. ****p<0.0001.

**Supplemental Figure 3. Standard chemotherapy induces DNA damage response programs in the *Dnmt3a*$^{R878H}$-mutated LSCs.** (A) Dose-response curve of cultured DNF and NF AML cells following 48 hours treatment with increasing concentrations of Dox+Arac, assessed by MTS assay. (B) Flow cytometry analysis of γH2AX phosphorylation in vehicle- and Dox+AraC-treated DNF and NF AML cells in culture following 48 hours of treatment. *In vivo* γH2AX levels in DNF and NF AML BM cells measured prior to treatment (T0) and 48 hours following chemotherapy (T48) in (C) bulk CD45.2$^{+}$ AML and (D) LSC-enriched LSK cells. (E) Heatmap of GSEA Hallmark NES scores for the indicated gene signatures in vehicle- and Dox+AraC-treated DNF and NF LSCs. Purple denotes negative enrichment and yellow denotes positive enrichment. Statistical significance from GSEA is indicated by asterisks, $^{\#}p<0.05$, *FDR<0.05, **FDR<0.01, ***FDR<0.001. (F) Frequency of G0 cells within DNF and NF LSK population. Statistical significance was determined using One-way (B) or Two-way ANOVA (C-E) or unpaired Welch's T-test (A, G) as indicated. **p<0.01 and ***p<0.001.

**Supplemental Figure 4. Single-cell gene transcriptional profiling of *Npm1^c^-Flt3^ITD^* LSK population reveals heterogeneity within the DNF and NF LSC compartments.** (A) Dot plot showing the percentage of cells with the marker gene expression within transcriptionally defined LSK clusters. (B) Heatmap of average cluster single cell GSEA scores of established stem and progenitor gene signatures (2-4). (C) UMAP visualization of WT LSK cells (n=1), colored by transcriptionally defined stem and progenitor compartments. Violin plot of single cell GSEA comparing (D) LSC-associated gene signature scores, and (E) quiescence signature scores in NF and DNF LSC-containing LSKs following vehicle or Dox+AraC treatment (n=1-2 pooled independent biological replicates per genotype). Dot and error bars represent mean ± SD, respectively. (F) Proportion of cells in G0-G1, S or G2 cell cycle phases within the NF and DNF HSC-like quiescent compartment after vehicle or Dox+AraC treatments.

**Supplemental Figure 5. Cell cycle independent, venetoclax plus azacitidine therapy reduces disease burden in *Dnmt3a$^{R878H}$*-mutant to a similar extent as in *Dnmt3a$^{WT}$-Npm1$^{c}$-Flt3$^{ITD}$* AML.** Frequency of (A&B) granulocytes ($Gr1^{+}/Mac1^{+}$) and monocytes ($Gr1^{-}/Mac1^{+}$), determined by flow cytometric analysis of NF and DNF $CD45.2^{+}$ AML BM cells following treatment with single agent venetoclax (DNF, n=12), azacitidine, Ven+Aza combination or vehicle control (NF, n=11; DNF, n=16 per treatment group unless otherwise specified). All four treatment arms were performed in parallel. Absolute number of (C) monocytes ($Gr1^{-}/Mac1^{+}$) and (D) granulocytes ($Gr1^{+}/Mac1^{+}$) in Ven+Aza-treated NF and DNF $CD45.2^{+}$ AML BM cells compared with vehicle controls. Absolute number of (E) circulating AML blasts in PB, (F) BM AML blasts, (G) AML monocytes, (H) LSC-enriched MPP3 and (I) MPP2 cells following single-agent venetoclax, azacitidine, or Ven+Aza treatment in NF and DNF AML compared with vehicle controls. (J) Survival analysis of mice transplanted with isolated MPP3 (n=3) or MPP2 (LSK, $CD48^{+}$, $CD150^{+}$; n=2) cells from DNF AML in a pilot study. Statistical significance was determined using the log-rank test. Each data point represents an independent biological replicate with median indicated by the horizontal line. Identical symbols indicate technical replicates derived from the same biological sample pooled across multiple independent experiments. Statistical significance was determined using Ordinary One-way ANOVA (A-B, E-I) or unpaired Mann Whitney test (C-D). *p<0.05, **p<0.01 and ****p<0.0001.

**Supplemental Figure 6. Chemotherapy effectively depletes *Dnmt3a$^{R878H}$*-mutated LSCs when combined with PegIFNα.** (A) Experimental design used to assess the effect of a single dose of pegIFNα in DNF AML. (B) Number of AML leukocytes in PB, (C) spleen weight, and (D) number of AML blasts in BM. (E) Absolute number of monocytes ($Gr1^{-}Mac1^{+}$), granulocytes ($Gr1^{+}Mac1^{+}$), and c-$Kit^{+}$ cells within the $CD45.2^{+}$ AML of the BM 48 hours post pegIFNα administration. Frequency and absolute number of (H) monocytes ($Gr1^{-}Mac1^{+}$), (I) granulocytes ($Gr1^{+}Mac1^{+}$), and (J) c-$Kit^{+}$ cells within $CD45.2^{+}$ BM AML cells from DNF mice. (K) Representative flow cytometry plot showing Sca-1 induction and gating strategy for LK, LSK, and combined c-$Kit^{+}$ populations within the HSPC compartment. Absolute number of (L) $Lin^{-}$c-$Kit^{+}$ cells (LK+LSK gates) and (M) MPP2 cells in DNF AML following Dox+AraC + pegIFNα compared with Dox+AraC alone. Biological replicates are shown with matching symbols indicating technical replicates derived from the same biological sample with median indicated by the horizontal line, combined across three independent experiments. Statistical significance was determined using an unpaired Mann Whitney test. #p=0.06, *p<0.05, **p<0.01, ****p<0.0001.

1. Vassiliou GS, Cooper JL, Rad R, Li J, Rice S, Uren A, et al. Mutant nucleophosmin and cooperating pathways drive leukemia initiation and progression in mice. Nat Genet. 2011;43(5):470-5.
2. Sommerkamp P, Romero-Mulero MC, Narr A, Ladel L, Hustin L, Schonberger K, et al. Mouse multipotent progenitor 5 cells are located at the interphase between hematopoietic stem and progenitor cells. Blood. 2021;137(23):3218-24.
3. Zeng AGX, Bansal S, Jin L, Mitchell A, Chen WC, Abbas HA, et al. A cellular hierarchy framework for understanding heterogeneity and predicting drug response in acute myeloid leukemia. Nat Med. 2022;28(6):1212-23.
4. Pietras EM, Reynaud D, Kang YA, Carlin D, Calero-Nieto FJ, Leavitt AD, et al. Functionally Distinct Subsets of Lineage-Biased Multipotent Progenitors Control Blood Production in Normal and Regenerative Conditions. Cell Stem Cell. 2015;17(1):35-46.
